# Disrupted tRNA modification leads to intestinal mitochondrial dysfunction and microbial dysbiosis

**DOI:** 10.1101/2025.11.27.690007

**Authors:** Di Ran, Yongguo Zhang, Yueqing An, Ying Hu, Yinglin Xia, Jun Sun

**Author notes:** Corresponding Author: Dr. Jun Sun, PhD, Professor, AAAS Fellow, FAPS, AGA Fellow, Division of Gastroenterology and Hepatology Department of Medicine, University of Illinois Chicago, 840 S Wood Street, MC716, Chicago, Illinois, 60612, USA. **Conflict of interest:** The authors declare no conflict of interest. The funders played no role in the study design, the collection, analyses, or interpretation of data, the writing of the manuscript, or the decision to publish the results.

## Abstract

**Background and aims:** Transfer RNA (tRNA) modifications determine translation fidelity and efficiency. It occurs through the action of specific enzymes that modify the nucleotides within the tRNA molecule. Our previous study demonstrated tRNA modopathies and altered queuine-related metabolites in inflammatory bowel diseases. Queuine tRNA-ribosyltransferase catalytic subunit 1 (QTRT1) and QTRT 2 co-localize in mitochondria and form a heterodimeric TGT participating in tRNA Queuosine (tRNA-Q) modification. Human body acquires Queuine/Vitamin Q from intestinal microbiota or from diet. However, the roles of tRNA-Q modifications in the maintenance of intestinal mitochondrial homeostasis and microbiome are still unclear.

**Methods:** We used publicly available human IBD datasets, QTRT1 knockout (KO) mice, QTRT1 intestinal epithelial conditional KO (QTRT1^ΔIEC^) mice, cultured cell lines with QTRT1-specific siRNA, and organoids from patients with IBD to investigate the mechanism of tRNA-Q modifications in intestinal mitochondrial homeostasis and therapeutic potential in anti-inflammation.

**Results:** In single cell RNA sequencing datasets of human IBD, we identified significant reduced intestinal epithelial QTRT1 expression in the patients with Crohn’s Disease. Using publicly available datasets, we identified significantly changes of Vitamin Q-associated bacteria in human IBD, compared to the healthy control. Qtrt1^-/-^ mice had significant reduction of Q-associated bacteria, e.g., *Bacteroides*. Alcian Blue and Mucin-2 staining revealed mucosal barrier damage and disrupted homeostasis, with reduced colonic cell proliferation. Intestinal tight junction integrity was impaired in QTRT1-KO mice, as evidenced by reduced ZO-1 and increased Claudin-10 expression. QTRT1^ΔIEC^ mice also showed dysbiosis and disrupted TJs. ATP synthesis was significantly decreased in the colon of QTRT1-KO mice, accompanied by severe mitochondrial dysfunction: reduced mitochondrial quality, Cytochrome-C release, and mitochondrial DNA (mtDNA) leakage. Mitochondrial dysfunction contributed to colonic cell death, as shown by elevated expressions of Cleaved Caspase-3 and Cleaved Caspase-1, increased BAX/Bcl-2 ratio, and positive TUNEL signals. Elevated CDC42, CD14, and CD4 levels in QTRT1-KO colon suggested mucosal immune activation and tissue repair responses. QTRT1-deficient CaCO2-BBE cells showed mitochondrial dysfunction. Cytochrome-C and mito-DNA release leading to cell death characterized by elevated expressions of Cleaved Caspase-3 and Caspase-1, increased BAX/Bcl-2 ratio, and higher apoptosis rate. Organoids isolated from patients with IBD showed reduced levels of QTRT1 and dysfunctional mitochondria. Restoring mitochondrial function leads to enhanced QTRT1.

**Conclusions:** These findings underscore the critical role of QTRT1/Q-tRNA modification in maintaining intestinal and microbial homeostasis. Mechanistically, QTRT1 loss impacts mitochondrial integrity and mucosal homeostasis. Our study highlights the novel roles of tRNA-Q modification in maintaining mucosal barriers and innate immunity in intestinal health.

**What is already known about this subject?:**

- Eukaryotes acquire queuine (q), also known as Vitamin Q, as a micronutrient factor from intestinal microbiota or from diet.
- Vitamin Q is needed for queuosine (Q) modification of tRNAs for the protein translation rate and fidelity.
- Queuine tRNA-ribosyltransferase catalytic subunit 1 (QTRT1) is reduced in human IBD.
- However, health consequences of disturbed availability of queuine and altered Q-tRNA modification in digestive diseases remain to be investigated.

**What are the new findings?:**

- QTRT1 deficiency leads to altered microbiome and reduced Vitamin Q-associated bacteria in human IBD and a QTRT1 KO animal model.
- QTRT1 protects the host against losing intestinal integrity during inflammation.
- QTRT1 localizes in mitochondria and plays novel functions by maintaining intestinal mitochondrial function. QTRT1 loss impacts tRNA modification in the intestine, linking to mitochondrial integrity and mucosal homeostasis.
- Human IBD showed reduced levels of QTRT1 and dysfunctional mitochondria. Restoring mitochondrial function leads to enhanced QTRT1.

**How might it impact on clinical practice in the foreseeable future?:** Targeting tRNA-Q modification in enhancing mitochondrial function will be a novel method to maintain intestinal health.

## Introduction

Eukaryotes acquire queuine, a micronutrient factor (Vitamin Q), from microbiota or diet. Vitamin Q incorporates into the wobble position of transfer RNA (tRNA) for Q-tRNA modification. tRNA modification affects fidelity and efficiency of translation from RNA to proteins. It occurs through the action of specific enzymes that modify the nucleotides within the tRNA molecule. Queuine tRNA-ribosyltransferase catalytic subunit 1 (QTRT1) and QTRT D co-localize in mitochondria and form a heterodimeric TGT participating in tRNA Queuosine (tRNA-Q) modification. Our previous study demonstrated tRNA modopathies and altered queuine-related metabolites in inflammatory bowel diseases ^1^. However, the roles of tRNA-Q modifications in the maintenance of intestinal homeostasis and innate immunity are still largely unclear.

Intestinal homeostasis is highly dependent on the proper functioning and distribution of microbiome and intestinal epithelial cells (IECs), the major component shaping the intestinal physiology. At the molecular level, many micronutrients are microbiome-dependent, they are not produced in the body and must be derived from the diet. Considering the dynamic nature of microbiome in human gastrointestinal tract (GI), the proportion of queuine absorbed from the microbiota dynamically regulates the disease progression ^2^. Q availability in a microbiome would affect microbial functionality, such as biofilm formation and virulence in bacteria ^3^. The synthesis of queuine by colonic gut microbiome is via cross-feeding based on a recent study ^4^. However, there is no study on Q-tRNA modification that may affect intestinal functions through alterations in gene expression. The gene expression differences in the host in turn help coordinate biosynthetic processes or population dynamics of microbiome. At the cellular level, our recent data indicate that the Q-tRNA modification not only depends on the gut microbiome but also affects cell proliferation and intestinal permeability through junction proteins ^1^. Interestingly, tRNA ribosyltransferase catalytic subunit 1&D (QTRT1/QTRTD) locate in the mitochondria ^5^. Given the critical role of mitochondria in energy production, redox balance, cell survival, microbiome, and inflammation ^6–8^, QTRT1 deficiency may lead to mitochondrial dysfunction, thus contributing to epithelial barrier disruption and inflammation. Investigations on the microbiome-derived tRNA modification and mitochondrial QTRT1 in the intestine will uncover novel mechanisms for mucosal disruption and inflammation.

In the current study, we hypothesize that tRNA-Q modifications by QTRT1 are essential for the maintenance of intestinal mitochondrial homeostasis and innate immunity. We used human IBD datasets, QTRT1 knockout (KO) mice, QTRT1^ΔIEC^ mice, cultured cell lines with QTRT1-specific siRNA, and organoids from patients with IBD to investigate the mechanism of tRNA-Q modifications in intestinal mitochondrial homeostasis and mucosal integrity. We explored the novel strategies by targeting QTRT1 level and mitochondria to restore intestinal health. Our study provides insights into the role of tRNA medication in the intestinal physiology and pathophysiology in inflammation.

## Results

### Downregulated QTRT1 expression and bacterial dysbiosis in IBD patients

To determine the clinical relevance of QTRT1 in human IBD, we reanalyzed single cell RNA sequencing datasets in the NCBI GEO database (accession GSE164985). We found the gene expression level of QTRT1 in epithelial cells was significantly downregulated in CD patients compared with healthy controls (**Fig.1A**). Human body acquires Vitamin Q from intestinal microbiota or from diet ^2^. Using publicly available datasets, we further identified significant changes of Vitamin Q-associated bacteria, e.g., *Bacteroides* and *Alistipes* were significantly reduced in IBD patients compared with healthy controls, based on analyses of 9 publicly available human gut metagenomic datasets **(Fig. 1B and 1C**).

**Fig.1.**
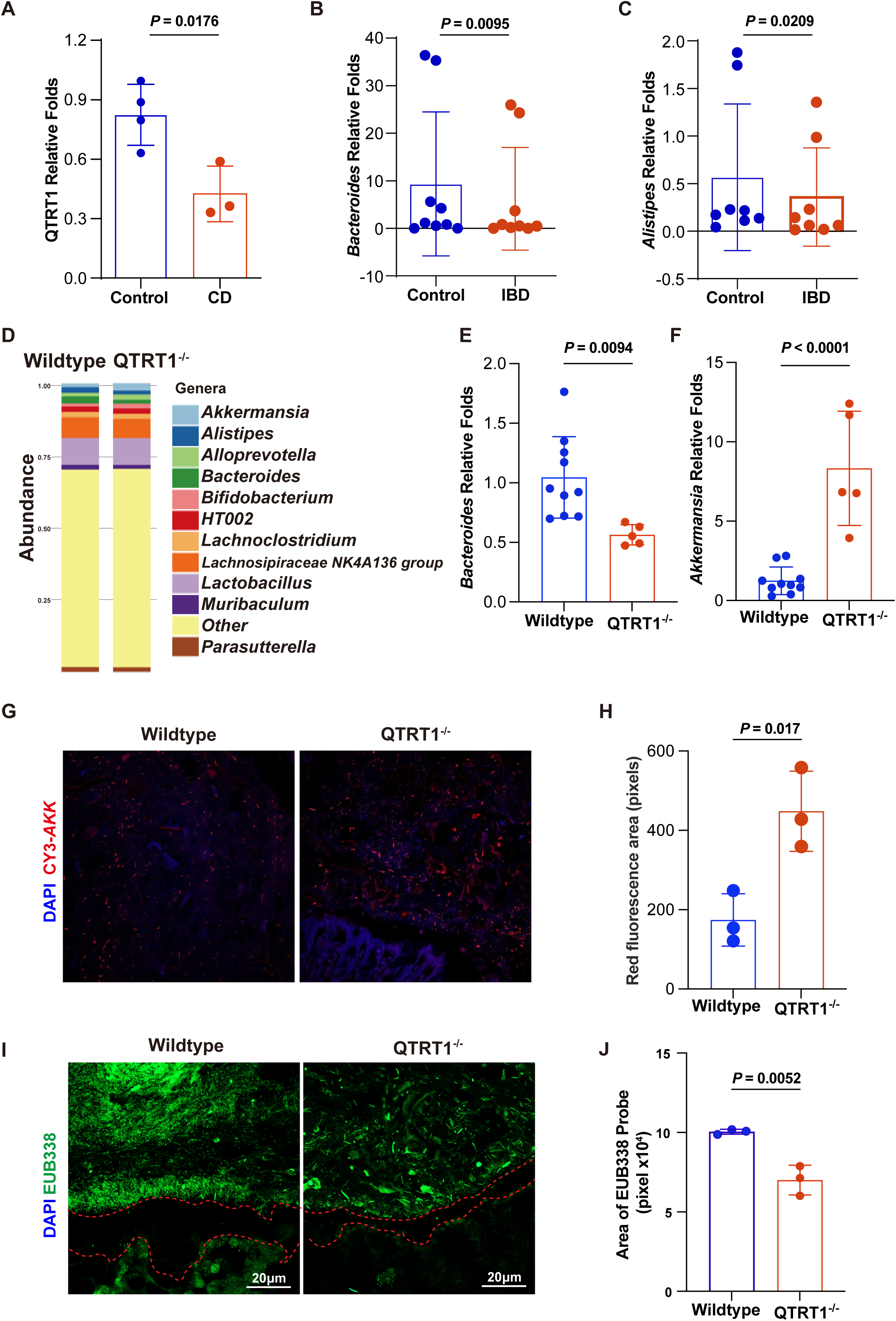
Reduced QTRT1 and altered Q-associated bacteria in human IBD and an animal model. **(A)** QTRT1 expression was significantly downregulated in colonic epithelial cells from patients with Crohn’s Disease, compared with healthy controls as determined by scRNA-seq. The data were retrieved from NCBI GEO scRNA-seq database (accession GSE164985), which included 43,692 cells from three CD patients and 51,036 cells from four healthy controls. Data are shown as mean ± SD, Welch’s t-test. **(B)** Q-associated bacteria *Bacteroides* (k=9 datasets) and **(C)** *Alistipe* (k=8 datasets) were significantly reduced in IBD patients compared with healthy controls at species level, based on analyses of publicly available human gut metagenomic datasets. Data are expressed as means ±SD, Wilcoxon rank sum test. **(D)** QTRT1^-/-^ mice exhibited 12 unique OTUs of microbiome, compared to the wild-type mice, based on 16s rRNA sequencing of the fecal samples. There are significant differences in bacteria: decreased *Bacteroides* and *Alistipe, and* increased *Akkermansia*, compared with Wildtype mice. (**E)** Decreased *Bacteroides* and (**F**) increased *Akkermansia* in QTRT1^-/-^ mice was determined by RT-PCR. Data are expressed as the mean□±□SD. Wildtype mice n=10, QTRT1^-/-^ mice n=5. Welch’s *t-*test. **(G)** *Akkermansia* in the colons of QTRT1^-/-^ and Wildtype mice was detected by fluorescence in situ FISH using the *Akkermansia*-specific Cy3-labeled probe (*AKK*-Cy3). **(H)** Quantification of the Cy3-positive area (in pixels) from FISH images. n□=□3 mice/genotype. Welch’s *t-*test. **(I)** Bacteria in the colon of QTRT1^-/-^ and Wildtype mice were found by fluorescence in situ hybridization. **(J)** Quantification of the EUB338-positive area (in pixels) from FISH images. n□=□3 mice/genotype. The red dashed line outlines the epithelial surface. The scale bar is 20□μm. All *P*-values are shown in the figures.

### QTRT1 deficiency leads to an altered microbiome

By measuring the 16sRNA DNA sequencing of the fecal samples, we identified exhibited 11 unique OTUs of microbiome compared to the WT mice: significant differences in bacterial genus, such as *Akkermansia*, *Alistipes*, *Bacteroides, Lactobacillus*, and *Muribaculaceae,* suggest potential physiological or metabolic impacts of QTRT1 on microbial ecology (**Fig. 1D**). Furthermore, QTRT1^-/-^ mice had significant reduction of Q-associated bacteria, e.g., *Bacteroides* **(Fig. 1D**). These sequencing findings were validated by qPCR, which showed decreased *Bacteroides* and increased *Akkermansia* abundance in QTRT1^-/-^ mice (**Fig. 1E–F**). Using species-specific FISH staining with CY3-*AKK* probe, we observed markedly increased *Akkermansia* in the colon of QTRT1^-/-^ compared to the WT mice (**Fig. 1G**), with quantification confirming significantly enlarged red fluorescence areas (**Fig. 1H**).

To understand the impact of QTRT1 on intestinal health, we investigated the four layers of the intestinal barrier: microbiome, mucin, epithelial cells, and local immune cells using QTRT1 KO and wildtype (WT) controls. We first assessed microbial localization and the integrity of the outer microbial layer. Universal bacterial FISH staining with EUB338 probe demonstrated an overall reduction in total microbial density in QTRT1^-/-^ mice (**Fig. 1I and 1J**).

### QTRT1 deficiency impairs Goblet cells and their mucin secretion in the intestinal mucosa

The FISH staining allows us to visualize the mucosal/chemical layer, the second barrier layer, in the colon (**Fig. 1I and 2A** indicated by the Red line). There was a significantly narrowed mucosal layer in the QTRT1^-/-^ mice, compared with the WT controls (**Figure 2B**). To investigate the impact of QTRT1 deficiency on intestinal barrier integrity, we used Alcian Blue staining to detect in Goblet cells. We found that the Goblet cells in the QTRT1-KO mice appeared smaller and contained less acidic mucins, with a substantial reduction in Goblet cells and mucin content in the colonic epithelium of QTRT1 mice, compared to WT controls (**Figure 2C**). Consistent with the change of Goblet cell number, MUC2 immunofluorescence staining further demonstrated a significant decrease in MUC2-positive Goblet cells in QTRT1-KO mice, compared to the Wildtype (WT) controls (**Figure 2D**).

**Fig. 2.**
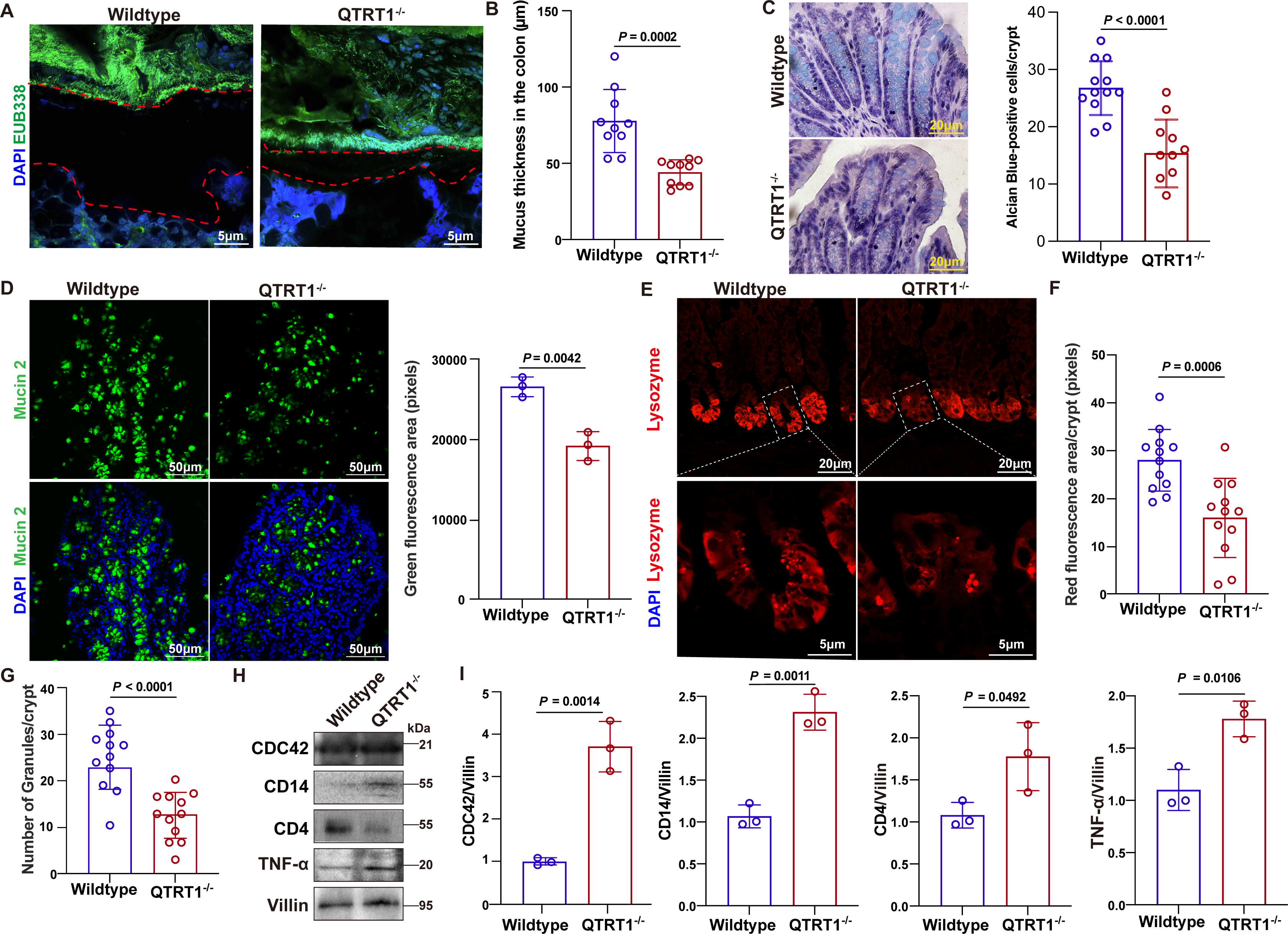
QTRT1 deficiency impairs Goblet cells and their mucin secretion, Paneth cell function and triggers immune activation in the intestinal mucosa. **(A)** Mucus thickness in the colon tissues of QTRT1^-/-^ and Wildtype mice was evaluated by FISH staining with EUB338 for all the bacterial species; the scale bar is 5□μm. **(B)** Quantification of mucus layer thickness in the colon. Three points were randomly selected for each mouse. n□=□10. **(C)** Representative images of Alcian Blue/PAS staining of the colon of QTRT1^-/-^ and Wildtype mice. The scale bar is 200□μm. Images are from a single experiment and represent three mice per group. Four crypts were randomly selected for each mouse. Data are expressed as mean ±SD., n = 12 for Wildtype, and n = 10 for QTRT1^-/-^, Welch’s *t-*test. **(D)** Representative confocal images of colonic tissues from wildtype and QTRT1^-/-^ mice stained for Mucin2 and DAPI. Data are expressed as mean ±SD, n = 3, Welch’s *t-*test. **(E)** Representative immunofluorescence staining of small intestinal sections from Wildtype and QTRT1^-/-^ mice. Sections were stained for Paneth cell marker Lysozyme (red). **(F)** Quantification of Lysozyme-positive cells per crypt shows a significant reduction in Paneth cell numbers in QTRT1^-/-^ mice. Four crypts were randomly selected for each mouse. Data are expressed as mean ±SD., n = 12, Welch’s *t-*test. **(G)** Number of lysozyme-positive granules per crypt demonstrating a significant decrease in Paneth cell granule content in QTRT1^-/-^ mice. Images are from a single experiment and represent three mice per group. Four crypts were randomly selected for each mouse. Data are expressed as mean ±SD., n = 12, Welch’s *t-*test. **(H and I)** Western Blot of colonic epithelial cell lysates from Wildtype and QTRT1^-/-^ mice, showing the expression of CDC42, CD14, CD4, and TNF-α. Villin was used as a loading control. Data are expressed as mean ± SD. n = 3 mice/genotype, Welch’s *t-*test. All *P*-values are shown in the figures.

### QTRT1 loss affects Paneth cell function and triggers immune activation

The third layer of the intestinal barriers is a single-cell epithelium consisting of Goblet cells, absorptive enterocytes, enteroendocrine cells, and Paneth cells. Among these cells, Paneth cells play a critical role in maintaining intestinal homeostasis and host-microbial interactions by secreting antimicrobial peptides (AMPs) ^9–12^. We found that lysosomes in Paneth cells showed a significant accumulation of lysosomal structures in QTRT1-KO mice (**Figure 2E and 2F**). Quantification of Lysozyme-positive cells per crypt shows a significant reduction in Paneth cell numbers and granules per crypt in QTRT1⁻/⁻ mice, compared to the WT mice (**Figure 2G**). In addition, Western blot data demonstrated increased expression of immune activation markers, including CDC42 (cytoskeletal remodeling in immune responses), CD14 (macrophage activation), and CD4 (T-cell infiltration) in the colonic mucosa of QTRT-KO mice, compared to the WT mice (**Figure 2H-2I**).

### QTRT1 deficiency disrupts tight junction integrity and increases intestinal permeability

Given that QTRT1 deficiency disrupts goblet cell function, impairs Paneth cell homeostasis, and triggers immune activation, we next examined its impact on the epithelial tight junctions,^13^ a crucial determinant of barrier function. The expression of TJ protein ZO-1 was markedly reduced in the QTRT-KO group, compared to WT controls (**Figure 3A**). Claudins include over 27 family members ^14^. We examined Claudin 2, 7, and 10. Claudin-2, a pore-forming tight junction protein associated with increased barrier permeability, was significantly upregulated in QTRT1-KO mice. Claudin-10 was increased in the QTRT1-KO mice as well. In contrast, the expression levels of Claudin-7 remained unchanged between QTRT1-KO mice and WT mice (**Figure 3A**). The densitometry analysis of these blots showed significant changes in ZO-1, Claudin-2, and 10 in the KO mice, compared to the WT mice (**Figure 3B**). We further studied the distribution of these TJ proteins. The discontinuous and fragmented staining pattern of ZO-1 in the colon of epithelial cells without QTRT1 suggested structural disorganization of tight junction complexes (**Figure 3C and 3D).** Claudin-7 was still stable in the absence of QTRT1 (**Figure 3E and 3F).** Interestingly, the colonic Claudin-10 was significantly upregulated at the apical side of colonic epithelial cells in QTRT1-KO mice (**Figure 3G and 3H)**. Functionally, these alterations were associated with a significant rise in FITC-dextran translocation into serum, indicating increased epithelial permeability in QTRT1^-/-^ mice (**Fig. 3I**).

**Fig. 3.**
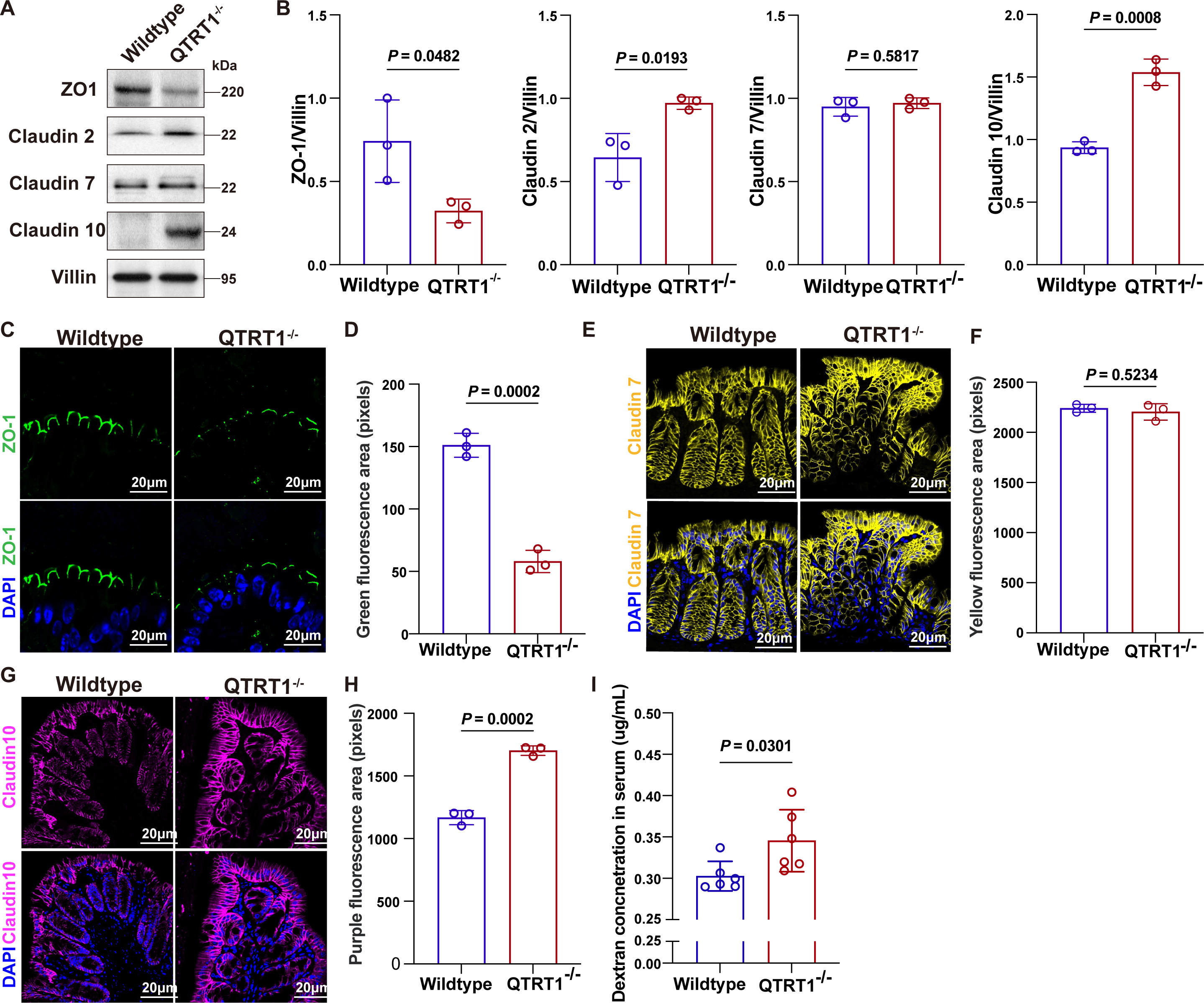
QTRT1 deficiency disrupts tight junction integrity and increases intestinal permeability. **(A, B)** Western blot analysis of tight junction proteins in colonic tissues from Wildtype and QTRT1^-/-^ mice. QTRT1 deficiency markedly decreased ZO-1 expression, while Claudin-2 and Claudin-10 levels were significantly increased. Claudin-7 remained unchanged. Villin was used as a loading control. Data are expressed as mean ± SD. n = 3 mice/genotype, Welch’s *t-*test. Immunofluorescence staining of **(C and D)** ZO-1 in colon tissues showed disrupted and discontinuous ZO-1 localization in QTRT1^-/-^ mice. **(E and F)** Claudin-7 (yellow) distribution remained intact in both wildtype and QTRT1^-/-^ mice, and **(G and H)** Claudin-10 (magenta) showed a prominent increase at the apical membrane of colonic epithelial cells in QTRT1^-/-^ mice. Nuclei are stained with DAPI (blue). The scale bar is 20□μm. All data are expressed as mean ± SD, Welch’s *t-*test, n = 3 mice/genotype. **(I)** QTRT1^-/-^ mice had significantly higher intestinal permeability levels compared to Wildtype mice. All data are expressed as mean ± SD, Welch’s *t-*test, n = 6 mice/genotype. All *P*-values are shown in the figures.

### QTRT1^Δ^^IEC^ mice exhibit impaired mucus barrier, reduced microbiome density, and disrupted epithelial junctions

To determine the intestinal epithelial role of QTRT1 in maintaining mucosal barrier homeostasis, we generated IEC-specific QTRT1 knockout mice, QTRT1^ΔIEC,^ by crossing QTRT1^LoxP^ mice with Villin-Cre drivers (**Fig. 4A**). Efficient loss of QTRT1 in intestinal epithelial cells was confirmed by immunoblotting (**Fig. 4B and 4C**). Loss of epithelial QTRT1 markedly reduced the thickness of the colonic mucus layer (**Fig. 4D and 4E**), accompanied by a significant decline in total mucosa-associated bacterial abundance, as quantified by EUB338 FISH staining (**Fig. 4D and 4F**). These findings indicate weakened mucus microbiota organization in QTRT1^ΔIEC^ mice. Immunofluorescence revealed a pronounced reduction and fragmentation of ZO-1 staining along epithelial junctions in QTRT1^ΔIEC^ colons (**Fig. 4G and 4H**), suggesting impaired tight-junction integrity. Consistently, ZO-1 protein levels were significantly decreased in QTRT1^ΔIEC^ colon detected by WB (**Fig. 4I and 4J**).

**Fig. 4.**
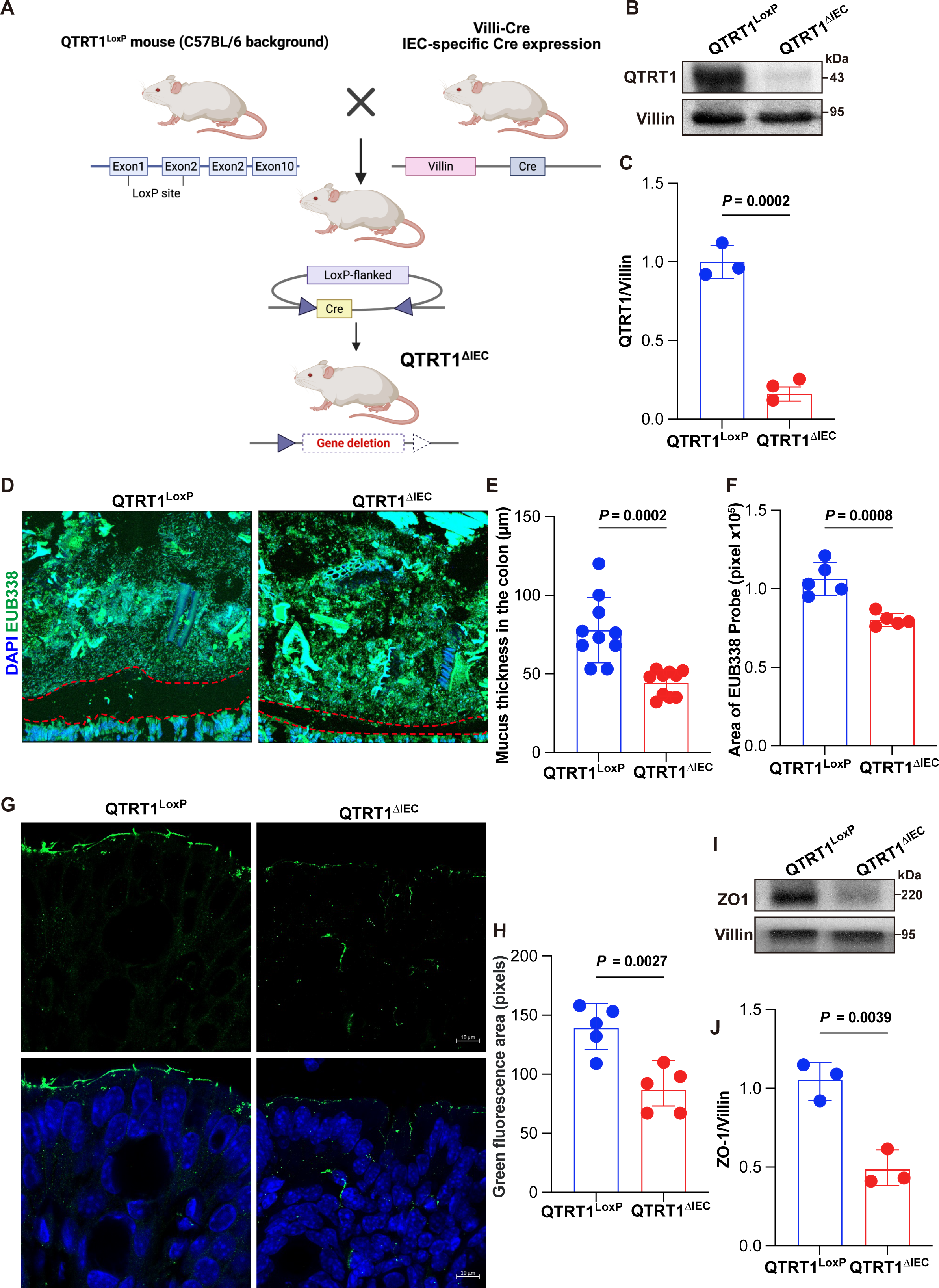
Characterization of QTRT1^ΔIEC^ mice and intestinal epithelial phenotype. **(A)** Schematic illustration of the generation of QTRT1^ΔIEC^ mice. QTRT1^LoxP^ mice (C57BL/6 background) were crossed with Villin-Cre mice to achieve intestine epithelial-specific deletion of QTRT1 in exons 1-3. **(B and C)** Western blot showed QTRT1 protein levels in isolated intestinal epithelial cells from QTRT1^LoxP^ and QTRT1^ΔIEC^ mice. Villin was used as a loading control. Data are expressed as mean ± SD. n = 3 mice/genotype, Welch’s *t-*test. **(D)** Quantification of the EUB338-positive area (in pixels) from FISH images. n□=□3 mice/genotype. The dashed red line denotes the mucus layer boundary. **(E)** Quantification of mucus thickness in colon sections. Data are expressed as mean ± SD. n = 10, Welch’s *t-*test. **(F)** Quantification of EUB338-positive bacterial area in colon sections. Data are expressed as mean ± SD. n = 5, Welch’s *t-*test. **(G and H)** Immunofluorescence staining of ZO-1 in colon tissues showed disrupted and discontinuous in QTRT1 ^ΔIEC^ mice. The scale bar is 20 µm. Data are expressed as mean ± SD. n = 5, Welch’s *t-*test. **(I and J)** Western Blot of ZO-1 in IECs from QTRT1^ΔIEC^ mice and QTRT1^LoxP^ mice, with Villin as a loading control. Data are expressed as mean ± SD. n = 3, Welch’s *t-*test. All *P*-values are shown in the figures.

### QTRT1 deficiency causes mitochondrial dysfunction and oxidative stress in the intestinal epithelium

Given the critical role of mitochondria in energy production, redox balance, cell survival, microbiome, and inflammation ^6–8^, we then examined whether QTRT1 regulates mitochondrial homeostasis in intestinal epithelial cells. We first assessed QTRT1’s subcellular localization. QTRT1 colocalized with the mitochondrial marker Tomm20 in WT colonic epithelium **(Fig. 5A).**

**Fig. 5.**
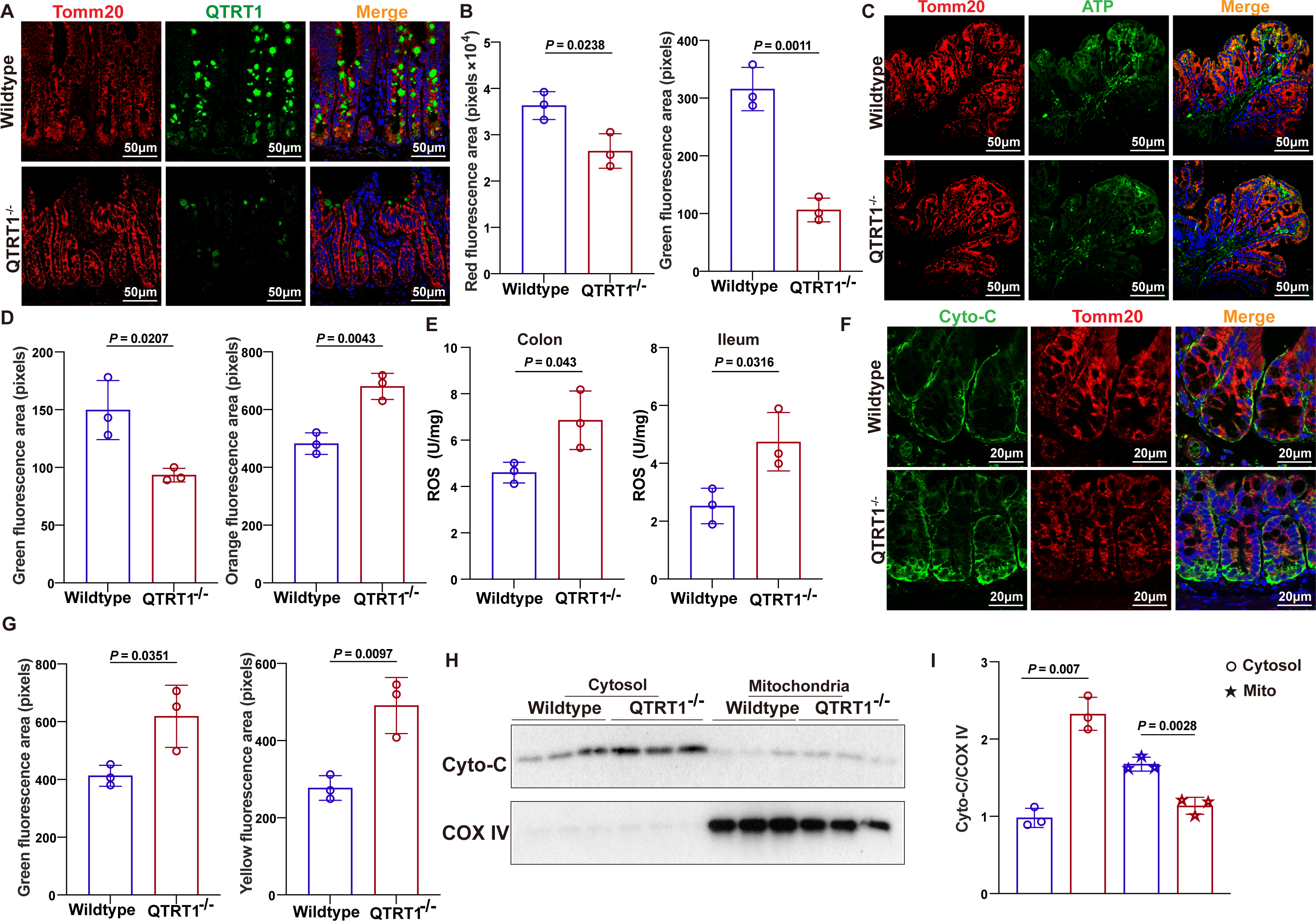
QTRT1 deficiency impairs mitochondrial function and elevates oxidative stress. **(A and B)** Immunofluorescence staining of colonic sections from Wildtype and QTRT1^-/-^ mice showing colocalization of QTRT1^-/-^ with the mitochondrial marker Tomm20 (red), QTRT1, and nuclei stained with DAPI (blue). The scale bar is 50□μm. Data are expressed as mean ± SD. n = 3 mice/genotype, Welch’s *t-*test. **(C and D)** ATP staining shows decreased mitochondrial ATP production in colonic sections of QTRT1^-/-^ mice compared to Wildtype controls. The scale bar is 50□μm. Data are expressed as mean ± SD. n = 3 mice/genotype, Welch’s *t-*test. **(E)** Quantification of reactive oxygen species (ROS) in colon and ileum tissue lysates from Wildtype and QTRT1^-/-^ mice. All data are expressed as mean ± SD, n = 3 mice/genotype, Welch’s *t-*test. **(F and G)** Immunofluorescence staining for Cytochrome C and Tomm20 (red) in colonic tissue. DAPI stains nuclei. The scale bar is 20□μm. All data are expressed as mean ± SD, n = 3 mice/genotype, Welch’s *t-*test. **(H and I)** Western blot analysis of Cytochrome C levels in cytosolic and mitochondrial fractions from colonic epithelial cells. COX IV serves as the mitochondrial loading control. All data are expressed as mean ± SD, n = 3 mice/genotype, Welch’s *t-*test. All *P*-values are shown in the figures.

QTRT1-KO mice exhibited a marked reduction of Tomm20 intensity, indicating decreased mitochondrial mass or structural integrity **(Fig. 5B**). Mitochondrial function was profoundly impaired in QTRT1 deficient intestines. ATP staining and quantification revealed significantly reduced ATP levels in the colonic mucosa of QTRT1^-/-^ mice (**Fig. 5C and 5D**). ROS assays demonstrated increased mitochondrial ROS accumulation in both the colon and ileum (**Fig. 5E**). To further evaluate mitochondrial stress, we examined cytochrome c distribution. QTRT1^-/-^colons showed increased cytosolic cytochrome c staining, together with altered Tomm20-defined mitochondrial morphology (**Fig. 5F and 5G**). WB of subcellular fractionation confirmed elevated cytochrome c in the cytosolic fraction and reduced retention within mitochondria (normalized to COX IV) in QTRT1^-/-^ tissues (**Fig. 5H and 5I**), indicating enhanced cytochrome c release and mitochondrial membrane stress in absence of QTRT1.

### QTRT1 deficiency promotes mtDNA release and induces apoptosis and pyroptosis, contributing to epithelial cell death

To determine whether mitochondrial damage in QTRT1-deficient intestines leads to cell death, we first assessed mitochondrial morphology and mtDNA distribution. WT epithelial cells displayed an intact Tomm20-defined mitochondrial network, whereas QTRT1^-/-^ mice exhibited fragmented and disorganized mitochondria accompanied by increased cytoplasmic and extracellular mtDNA signals (**Fig. 6A**). We then evaluated epithelial apoptosis. QTRT1^-/-^ colons showed a marked increase in cleaved-caspase-3 positive cells (**Fig. 6B**). Immunoblotting further demonstrated elevated cleaved-caspase-3, increased p-PARP, and upregulated pro-apoptotic Bax with concomitant reduction of anti-apoptotic BCL-2 (**Fig. 6C and 6D).** In addition to apoptosis, cleaved-caspase-1 levels (P20/P22) were significantly elevated in QTRT1^-/-^ mice (**Fig. 6C and 6D**), consistent with inflammasome activation and pyroptotic signaling. TUNEL staining further confirmed widespread epithelial cell death in QTRT1^-/-^ colons (**Fig. 6E and 6F**). Finally, assessment of epithelial proliferation showed a significant reduction in PCNA-positive cells in the QTRT1^-/-^ crypts (**Fig. 6G and 6H**).

**Fig. 6.**
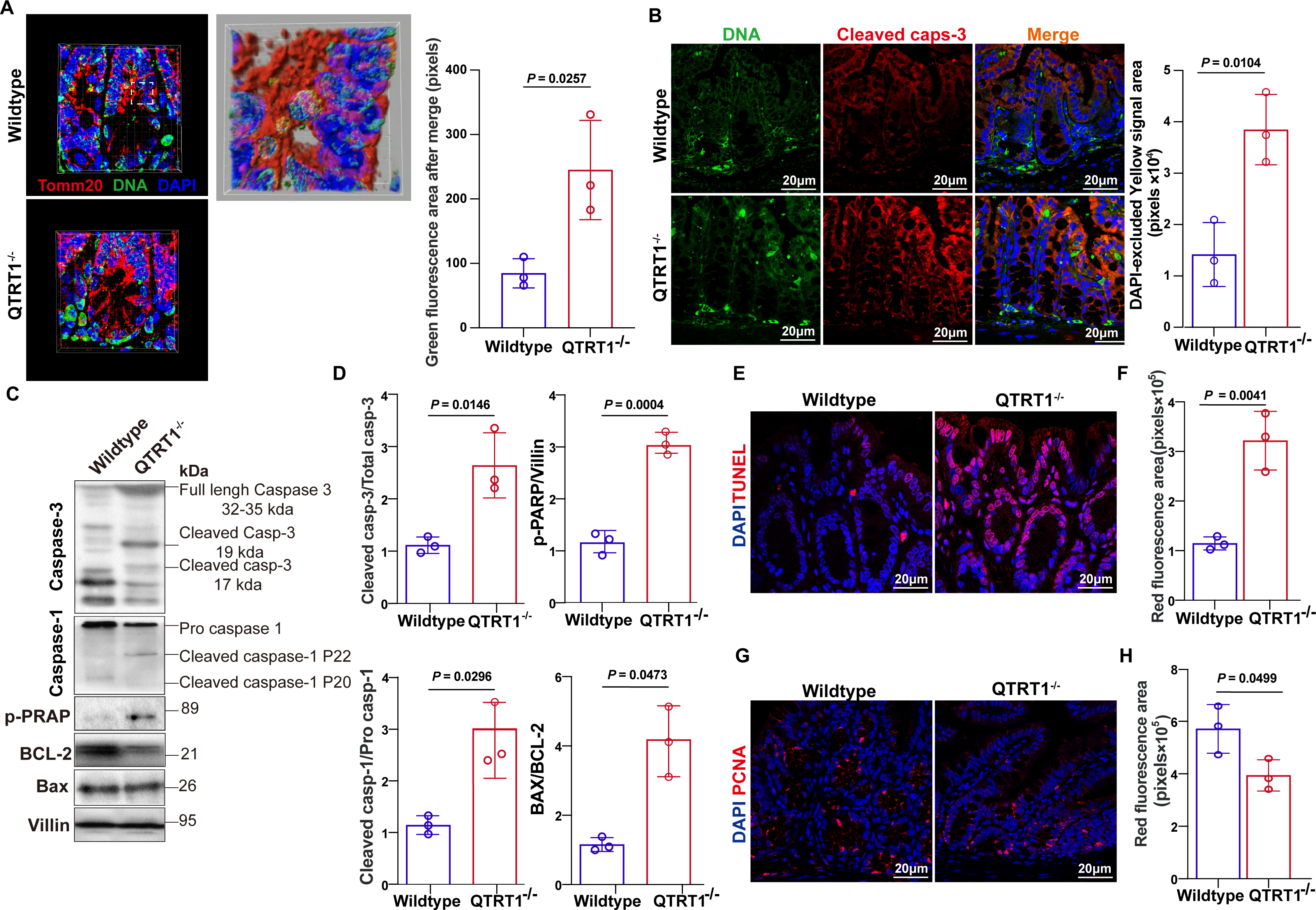
QTRT1 deficiency promotes mtDNA release and induces apoptosis and pyroptosis, contributing to epithelial cell death. **(A)** QTRT1^-/-^ mice display disrupted and fragmented mitochondrial networks along with increased cytoplasmic and extracellular mtDNA release by Immunofluorescence staining of colonic sections from Wildtype and QTRT1^-/-^ mice using Tomm20 (red) and mtDNA antibodies. All data are expressed as mean ± SD, n = 3 mice/genotype, Welch’s *t-*test. **(B)** Increased Cleaved Caspase-3 expression and DNA in the colonic epithelium of QTRT1^-/-^ mice by immunofluorescence staining. The scale bar is 20□μm. All data are expressed as mean ± SD, n = 3 mice/genotype, Welch’s *t-*test. **(C, D)** Western blot analysis of colon tissue lysates of increased Cleaved caspase-3, Bax, phosphorylated PARP (p-PARP), decreased expression of BCL-2, and increased BAX/BCL-2 in QTRT1^-/-^ mice. And the increase in Cleaved caspase-1 (P20/P22) levels. All data are expressed as mean ± SD, n = 3 mice/genotype, Welch’s *t-*test. **(E and F)** A marked increase in TUNEL-positive cells in QTRT1^-/-^ mice compared to wild-type controls by TUNEL staining of colon sections. The scale bar is 20□μm. All data are expressed as mean ± SD, n = 3 mice/genotype, Welch’s *t-*test. **(G and H)** The numbers of proliferating epithelial cells were observed by immunofluorescence staining of PCNA in colonic crypts. Nuclei are stained with DAPI (blue). The scale bar is 20□μm. All data are expressed as mean ± SD, n = 3 mice/genotype, Welch’s *t-*test. All *P*-values are shown in the figures.

### QTRT1 knockdown disrupts mitochondrial homeostasis by reducing ATP production and increasing oxidative stress *in vitro*

To investigate QTRT1’s direct regulation on mitochondrial function in epithelial cells, we generated QTRT1 KD using human CaCO-2 cells. Western blotting confirmed efficient depletion of QTRT1 protein (**Fig. 7A**). Loss of QTRT1 resulted in marked mitochondrial fragmentation and loss of the filamentous network defined by Tomm20 staining (**Fig. 7B-7D**), indicating structural instability of the mitochondrial compartment. Mitochondrial function was substantially impaired in QTRT1 KD cells. ATP fluorescence intensity was significantly reduced in knockdown cells (**Fig. 7D and 7E**), consistent with compromised bioenergetic capacity. MitoSOX staining revealed a pronounced increase in mitochondrial ROS in QTRT1-depleted cells **(Fig. 7F)**, demonstrating heightened oxidative stress. To further assess mitochondrial fitness, we evaluated mitochondrial membrane potential (MMP) using JC-1 assays (**Fig. 7G)**. QTRT1 knockdown cells exhibited a significant decrease in JC-1 red positive cells (**Fig. 7H)**, whereas MMP showed an increased trend in the QTRT1 knockdown cells (**Fig. 7I)**.

**Fig. 7.**
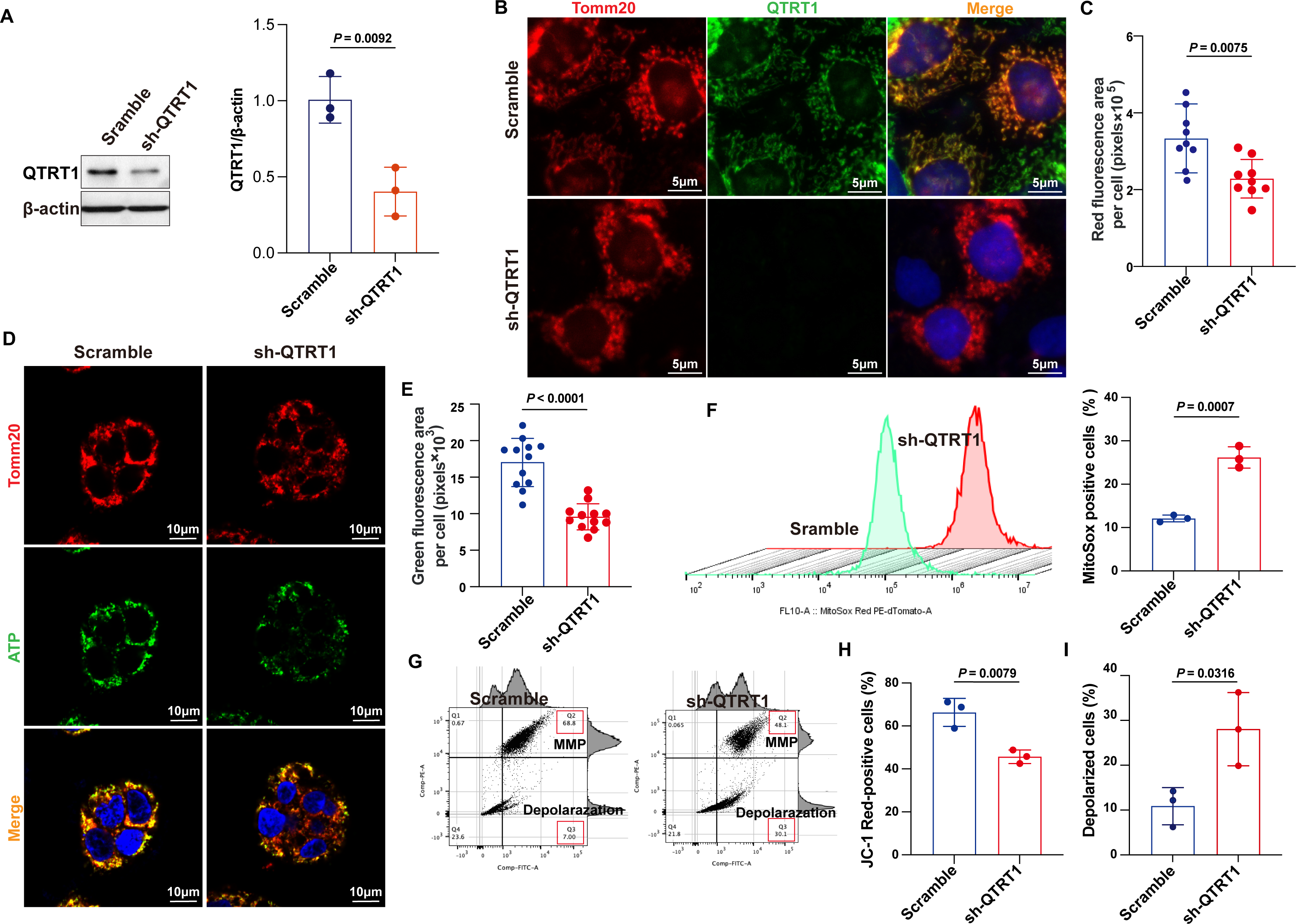
QTRT1 knockdown disrupts mitochondrial homeostasis by reducing ATP production and increasing oxidative stress *in vitro*. **(A)** Western blot confirming reduced QTRT1 protein levels in QTRT1-KD CaCO-2 cells compared to control. All data are expressed as mean ± SD, n = 3 independent experiments, Welch’s *t*-test. **(B and C)** Mitotracker-red staining shows fragmented and disorganized mitochondrial networks in QTRT1 KD cells. The scale bar is 5□μm. All data are expressed as mean ± SD, n = 3 individual experiments, each performed in triplicate. Welch’s *t*-test. **(D and E)** ATP generation from mitochondria stained by Tomm20 (red) was tested by immunofluorescence staining in QTRT1-KD CaCO-2 cells and *Scramble* CaCO-2 cells controls. The scale bar is 10 μm. All data are expressed as mean ± SD, n = 4 individual experiments, each performed in triplicate. Welch’s *t*-test. **(F)** MitoSox-Red staining reveals elevated mitochondrial ROS levels in QTRT1-KD cells by Flow cytometry. All data are expressed as mean ± SD, n = 3 individual experiments. Welch’s *t*-test. **(G, H, and I)** Flow cytometry was used to investigate mitochondrial membrane potential (MMP) and depolarized cells with JC-10 assays. All data are expressed as mean ± SD, n = 3 individual experiments. Welch’s *t*-test. All *P*-values are shown in the figures.

### QTRT1 knockdown triggers mitochondrial apoptosis, pyroptosis, and compensatory mitophagy *in vitro*

To further define the cellular consequences of QTRT1 depletion, we examined mitochondrial integrity and apoptotic pathways in the QTRT1 KD CaCO-2 cells. Loss of QTRT1 resulted in a marked reduction of the mitochondrial TOMM20 and a significant increase in cytosolic Cytochrome C with a corresponding decrease in mitochondrial Cytochrome C (**Fig. 8A and 8B**). QTRT1 KD also activated apoptotic and pyroptotic pathways, as evidenced by increased Cleaved caspase-3, elevated Bax/Bcl-2 ratio, and enhanced Cleaved caspase-1 levels (**Fig. 8A and 8B**). Flow cytometry confirmed a significantly higher proportion of apoptotic cells in QTRT1 depleted cultures (**Fig. 8C and 8D**). Given the extent of mitochondrial damage, we next investigated whether mitophagy was engaged as a compensatory mechanism. Mitophagy dye and lysosomal colocalization assays demonstrated enhanced mitophagy flux in the QTRT1 KD cells (**Fig. 8E-8G**).

**Fig. 8.**
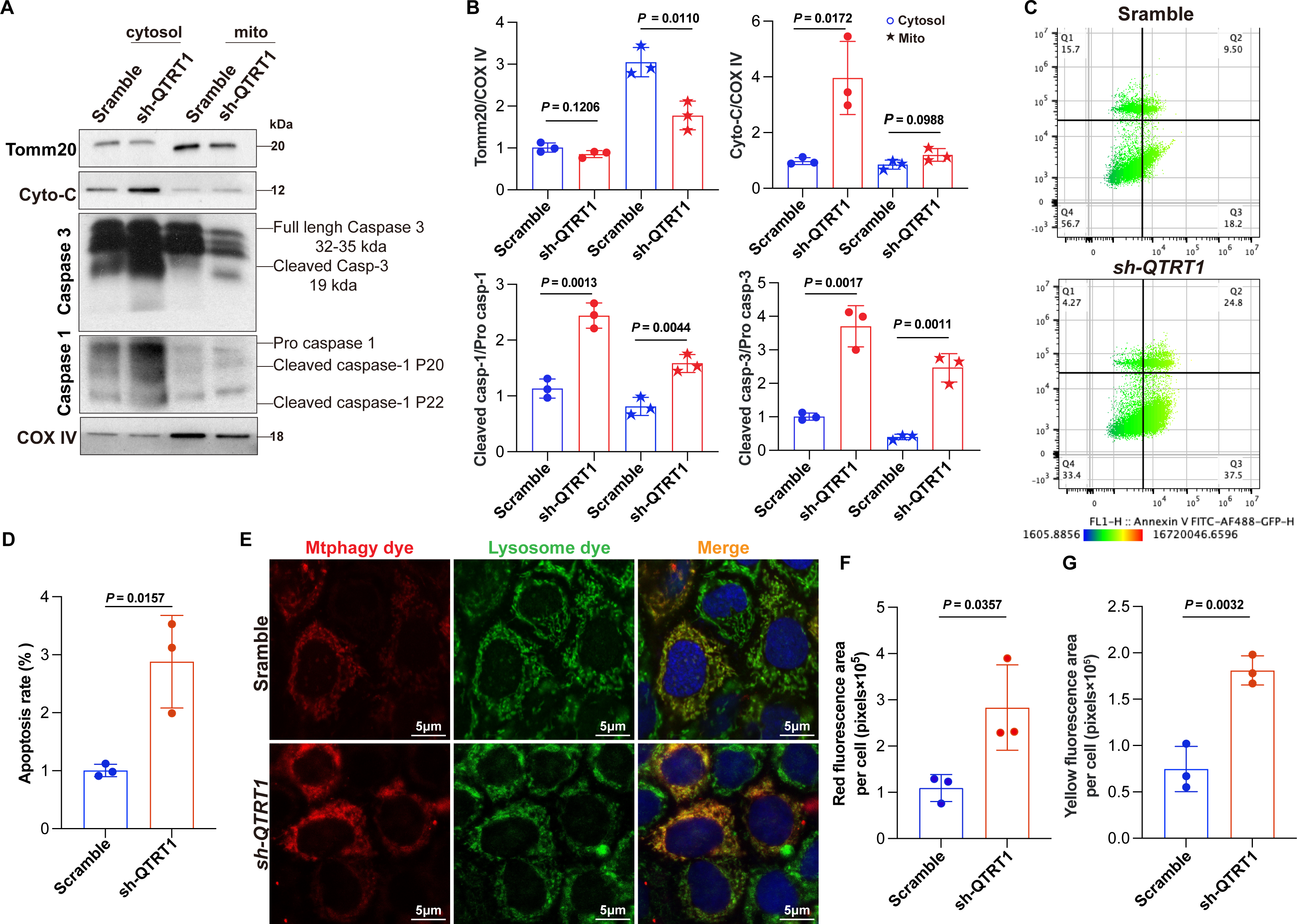
QTRT1 knockdown in CaCO2-BBE cells disrupts mitochondrial homeostasis. **(A and B)** Decreased expression of mitochondrial markers Tomm20 and mitochondrial Cytochrome C, accompanied by increased cytoplasmic Cytochrome C, in QTRT1 KD CaCO-2 cells was tested by Western Blot. And upregulation of Cleaved Caspase-3 and Cleaved Caspase-1 was detected by Western blot analysis. All data are expressed as mean ± SD, n = 3 individual experiments. Welch’s *t*-test. **(C and D)** Flow cytometry analysis reveals a higher proportion of apoptotic cells in QTRT1 KD compared to control cells. All data are expressed as mean ± SD, n = 3 individual experiments. Welch’s *t*-test. **(E-G)** Mtphagy dye and lysosome dye co-staining show increased mitophagy activity in QTRT1-KD cells. The scale bar is 5□μm. All data are expressed as mean ± SD, n = 3 individual experiments. Welch’s *t*-test. All *P*-values are shown in the figures.

### Reduced QTRT1 and mitochondrial dysfunction in the IBD organoids

To determine impacts of QTRT1 on the intestinal epithelium, we generated patient derived organoids using colonoscopic biopsies from healthy controls and individuals with IBD (**Fig. 9A**). Immunofluorescence staining showed significantly lower QTRT1 expression in the IBD organoids (**Fig. 9B and 9C**). This reduction coincided with the decreased mitochondrial marker Tomm20 (**Fig. 9B and 9D**). Using a MitoTracker Red live imaging (**Fig. 9E**), we were able to observe the dynamic changes of mitochondria. There was markedly reduced mitochondrial signal intensity in IBD organoids, compared to alive normal organoids (**Fig. 9F).** The red dye per crypts was also significantly reduced in the IBD alive organoids **(Fig. 9G**).

**Figure 9.**
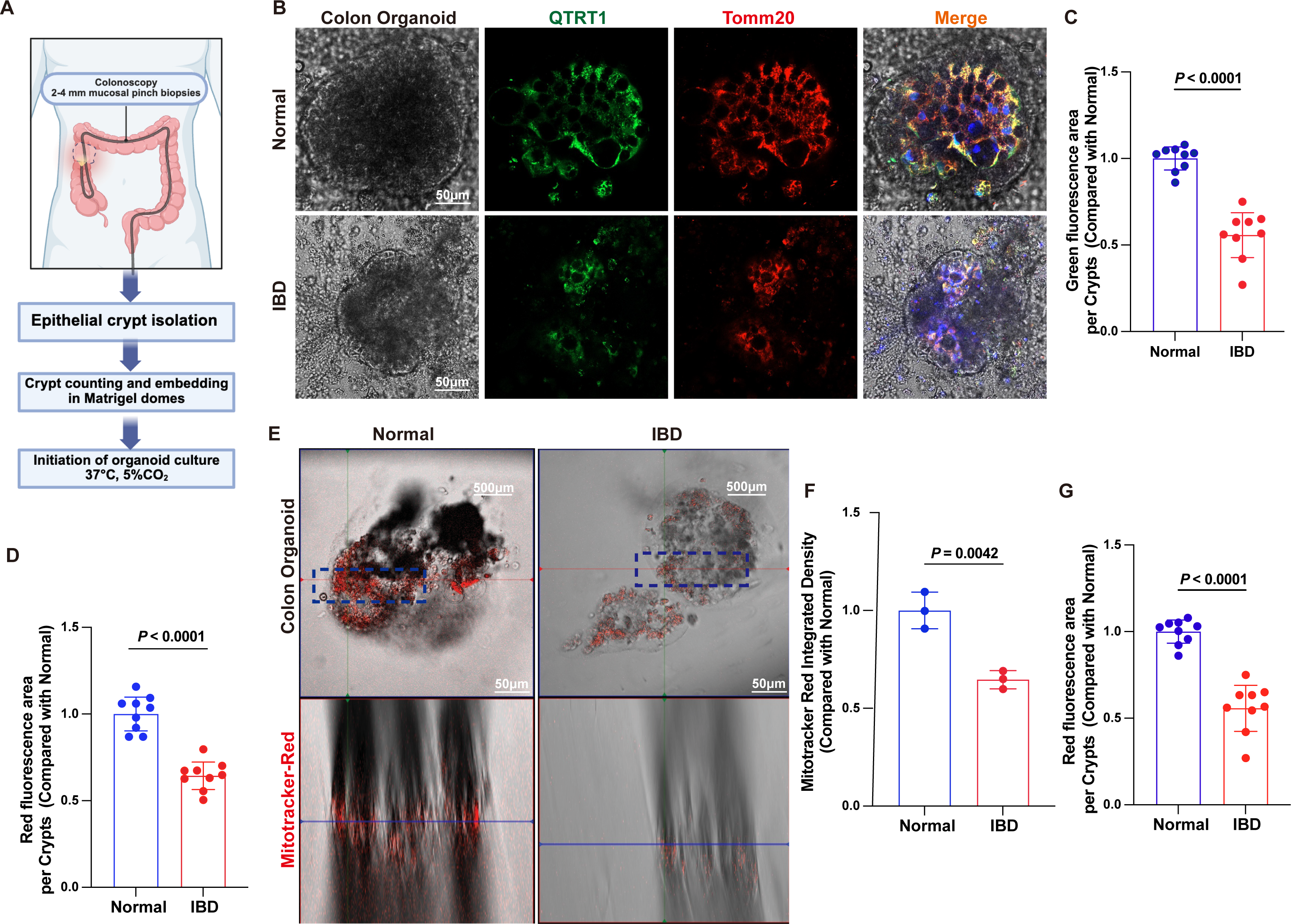
Reduced QTRT1 expression and mitochondrial dysfunction in IBD colon organoids. **(A)** Schematic overview of human colon organoid generation. **(B)** Representative immunofluorescence staining of colon organoids with Tomm20 and QTRT1. (**C**) QTRT1 (green) and **(D)** Mitochondrial marker Tomm20 (red) fluorescence area per crypt, significant reduced in epithelial in IBD patient organoids. The scale bar is 50□μm. All data are expressed as mean ± SD, n = 3 patients. Three crypts were randomly selected for each organoid. Welch’s *t*-test. **(E)** Representative brightfield and Mitotracker-red imaging of colon organoids derived from health control and IBD patients. All data are expressed as mean ± SD, n = 3 patients, Welch’s *t*-test. **(F and G)** Quantification of Mitotracker-red integrated density and red fluorescence area per crypt. All data are expressed as mean ± SD, n = 3 patients. In Figure **G**, three crypts were randomly selected for each organoid. Welch’s *t*-test. All *P*-values are shown in the figures.

### Restored the QTRT1 deficiency and mitochondrial by MitoQ in organoids from human IBD

We then tried to rescue mitochondrial defects in human IBD epithelium by treating human organoids with 10 μM MitoQ for 24 hours (**Fig. 10A**). IBD organoids without MitoQ treatment exhibited markedly reduced Tomm20 and QTRT1 fluorescence compared with healthy controls. Mito-Q treatment significantly restored Tomm20 intensity (**Fig. 10A and 10B**). While Mito-Q had minimal effects on normal organoids, it markedly increased QTRT1 in IBD organoids (**Fig. 10A and 10C**).

**Figure 10.**
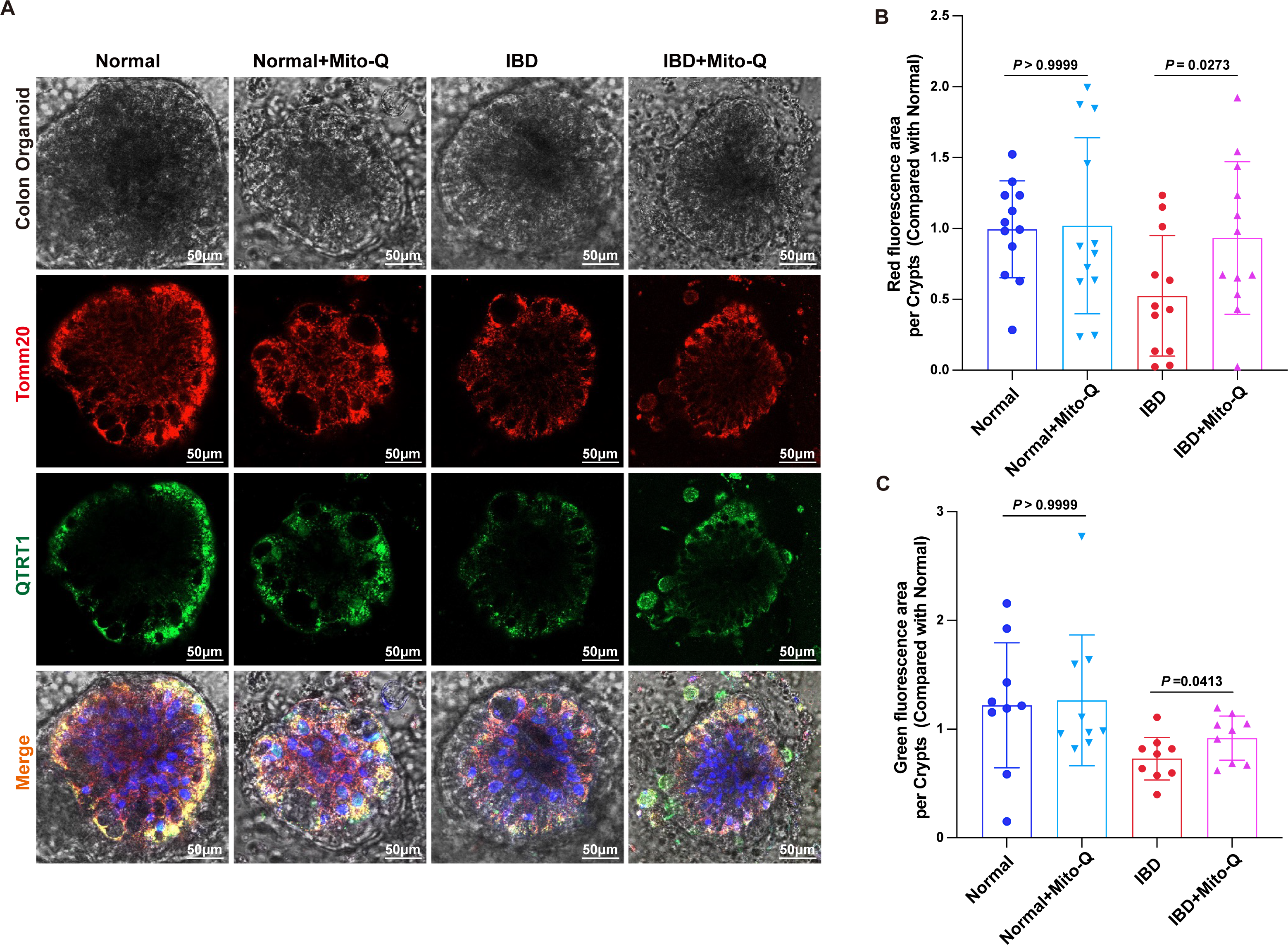
Mito-Q treatment restores mitochondrial and QTRT1 expression in IBD colon organoids. **(A).** 10 μM Mito-Q treatment of normal and IBD colon organoids for 24 h. Representative immunofluorescence images of colon organoids labeled for QTRT1 and Tomm20 (red), with merged images including nuclei (blue). **(B)** Quantification of Tomm20 red fluorescence area and **(C)** QTRT1 green fluorescence per crypt. The scale bar is 50□μm. All data are expressed as mean ± SD, the data points present 3-4 organoids for each patient. Quantification was performed using three crypts per patient, with three patients per group, n = 3 patients. Welch’s ANOVA test was used in (B) and (C), followed by a post-hoc test using the Dunnett’s T3 multiple comparation test. All *P*-values are shown in the figures.

## Discussion

In the current study, we report a QTRT1 deficiency in the human IBD. Mechanistically, deletion of QTRT1 leads to dysbiosis through Q-associated bacteria, impairs intestinal barrier integrity, and induces mitochondrial dysfunction. Therapeutically, Mito-Q treatment in the organoids from patients with IBD is able to increase QTRT1 and restore mitochondrial integrity. Our current findings underscore the critical role of QTRT1/Q-tRNA modification in maintaining intestinal mucosal integrity and microbial homeostasis through mitochondrial function.

The intestinal mucosal integrity is maintained by microbial, chemical, cellular, and immunological layers. Specifically, these four barrier layers include an outer microbial layer containing gut microbiome, a mucus layer, a single-cell epithelial layer, and an inner lamina propria layer housing innate and adaptive immune cells. We identified the impacts of QTRT1 on all layers, suggesting QTRT1 plays critical role in the fundamental functions in the intestinal health. Vitamin Q is an elusive and lesser-recognized micronutrient. We found that Vitamin Q-associated bacteria were reduced in human IBD and also in the QTRT1 KO mice. Although Q-associated research in IBD is in its infancy stage, there is a report on IBD patients with significant changes of Q-associated bacterial profile ^3^. Intestinal dysbiosis may be improved by probiotic *Lactobacillus* sp. or *Bifidobacterium* sp, which are Q-associated bacteria ^3^. We found that Paneth cells a significant accumulation of lysosomal structures in QTRT1-KO mice, which may lead to disrupted antimicrobial peptide secretion and altered microbial defense mechanisms, and this aberrant lysosomal accumulation indicates compromised antimicrobial defense, potentially exacerbating microbial dysbiosis and barrier dysfunction. These findings suggest that QTRT1 plays a crucial role in Paneth cell homeostasis and immune regulation. QTRT1 deficiency also impairs goblet cell function, disrupts mucin secretion, and compromises microbial segregation within the intestinal mucosa.

QTRT1 status might not only impact the microbiome, but also affect intestinal permeability. Intestinal tight junction integrity was impaired in QTRT1-KO mice, as evidenced by reduced ZO-1 and increased Claudin-2 expression. Elevated CDC42, CD14, and CD4 levels in QTRT1-KO colon suggested mucosal immune activation and tissue repair responses. These findings of altered TJs align with our previous observations of mucus layer depletion, microbial encroachment, and immune activation in digestive diseases ^17, 18^. Thus, our study highlights the novel roles of tRNA-Q modification in maintaining mucosal barriers and innate immunity in intestinal health. Interestingly, mitochondrial QTRT1 was reported by Boland et al. ^5^. However, the mitochondrial QTRT1 impacts on the intestinal functions of has not been studied. We have demonstrated that ATP synthesis was significantly decreased in the colon of QTRT1-KO mice, accompanied by severe mitochondrial dysfunction: reduced mitochondrial quality, Cytochrome-C release, and mtDNA leakage. Mitochondrial dysfunction contributed to colonic cell death, as shown by elevated expressions of Cleaved Caspase-3 and Cleaved Caspase-1, increased BAX/Bcl-2 ratio, and positive TUNEL signals. Because ATP is essential for tight junction assembly, mucus secretion, and epithelial repair, this energy deficit may contribute to the observed intestinal barrier dysfunction. QTRT1-deficient CaCO2-BBE cells showed mitochondrial dysfunction, including reduced ATP synthesis, increased mitochondrial ROS generation, decreased mitochondrial membrane potential, and enhanced mitophagy. Cytochrome-C and mito-DNA release leading to cell death characterized by elevated expressions of Cleaved Caspase-3 and Caspase-1, increased BAX/Bcl-2 ratio, and higher apoptosis rate. Our in vitro findings confirm that QTRT1 depletion leads to mitochondrial dysfunction, characterized by increased mitochondrial fragmentation, reduced ATP production, elevated oxidative stress, impaired mitochondrial membrane potential, and enhanced mtDNA release. The consistency between our in vivo and in vitro results highlights QTRT1’s essential role in mitochondrial homeostasis and epithelial cell metabolism. QTRT1 is crucial for mitochondrial stability and epithelial cell survival. While mitophagy initially acts as a protective response, excessive mitochondrial degradation may contribute to cellular energy depletion, increased apoptosis, and impaired epithelial regeneration, further emphasizing QTRT1 essential role in intestinal homeostasis. It suggests that QTRT1 loss contributes to intestinal barrier dysfunction via mitochondrial impairment.

The etiology of IBD has been characterized as chronic intestinal inflammation resulting from many factors, including micronutrients and epigenetic mechanisms via DNA methylation and non-coding RNA^19^. Investigating human biopsies and reanalyzing human datasets, we show provocative data that QTRT1 is significantly downregulated in human IBD. QTRT1 deficiency leads to mitochondrial dysfunction, thus contributing to epithelial barrier disruption and inflammation. Our study has further demonstrated the reduced QTRT1 in IBD organoids is restored by MitQ through the mitochondrial functions. The supply of micronutrient Q can be altered in situations, such as during disease progression or when certain food types are limited. Changes in Q availability in a microbiome would affect its functionality, such as biofilm formation and virulence ^3^. However, mechanisms of how vitamin Q and Q-tRNA modifications regulate Q-producing microbiome in IBD are unknown ^15^. Q produced by microbiota is taken up from the colonic lumen into enterocytes through specific cellular uptake mechanisms followed by incorporation into the wobble anticodon position of the four tRNAs by a heterodimeric enzyme encoded in the human genome ^16^. We speculate that microbiome-dependent vitamin Q/Q-tRNA modification improves mucosal barriers and maintains intestinal homeostasis, and targeting Q-tRNA modopathies to protect host against chronic inflammation. The limited literature and merging interests in Q-tRNA in human diseases highlight the needs to work on t-RNA modification in the future.

In summary, we have demonstrated that QTRT1 deficiency leads to altered microbiome and reduced Vitamin Q-associated bacteria in human IBD and a QTRT1 KO animal model. QTRT1 protects the host against losing intestinal and microbial integrity during inflammation. QTRT1 localizes in mitochondria and plays novel functions by maintaining intestinal mitochondrial function. Restoring mitochondrial function leads to enhanced QTRT1 in organoids from human IBD. Thus, investigations on the intestinal microbiome-derived tRNA modification and mitochondrial QTRT1 will uncover novel mechanisms for mucosal disruption and chronic inflammation. This knowledge could be exploited to define novel strategies to prevent and treat IBD and other digestive diseases.

## Materials and Methods

### Meta-analysis of human IBD scRNA studies and human gut microbiota metagenomic studies

To investigate the clinical significance of QTRT1 in IBD, with a primary focus on Crohn’s disease, we conducted a comprehensive meta-analysis of publicly available IBD-related single-cell RNA sequencing (scRNA-seq) data from the NCBI Gene Expression Omnibus database (https://www.ncbi.nlm.nih.gov/geo/). Among these datasets, GSE164985 was identified as a key resource containing colonic epithelial biopsy samples from 3 CD patients and 4 healthy controls, enabling evaluation of QTRT1 expression within disease-affected epithelial compartments ^20^. Publicly available scRNA-seq datasets from IBD studies were processed through a standardized pipeline ^21^. QTRT1 expression values were extracted, normalized according to the processing pipeline required for each dataset, and compared between disease and control groups.

In addition to genomic profiling, we integrated publicly available human gut microbiota metagenomic datasets to explore potential relationships between microbial community composition and QTRT1 expression. Meta-analysis was using nine human gut microbiota metagenomic datasets of IBD patients compared with healthy controls ^22–30^. The analysis of bacterial relative abundance variations between IBD patients and healthy controls were calculated at species level.

### Isolation and culture of human colonic organoids

This study was conducted under approval from the University of Illinois Chicago Institutional Review Board (STUDY2023-0327). Human endoscopic biopsy specimens were obtained from patients at the University of Illinois Chicago Hospital for the establishment of human intestinal organoid cultures. Colonic crypts were isolated from the biopsy samples using a previously described protocol with minor modifications. Briefly, biopsies were washed in cold DPBS (without Ca²⁺/Mg²⁺) and incubated for 5 min to allow debris to settle. The supernatant was removed, and tissues were transferred to a 1.5 mL tube, minced thoroughly, and rinsed into a 15 mL tube with D-PBS. After gravity settling, the supernatant was removed, and samples were incubated with Gentle Cell Dissociation Reagent (STEMCELL Technologies) for 30 min at room temperature on a rocking platform. Tissues were washed (290g, 5 min), resuspended in ice-cold DMEM/F-12 containing 1% BSA, and vigorously pipetted to release crypts.

The suspension was passed through a 70 µm strainer, rinsed with additional DMEM/F-12 with1% BSA, and centrifuged at 290g for 5 min. The crypt pellet was resuspended in growth factor-reduced Matrigel and plated as 50 µL domes in pre-warmed 24-well plates. After polymerization at 37°C for 20 minutes, the domes were overlaid with IntestiCult Human Organoid Growth Medium (STEMCELL Technologies) and maintained at 37°C with 5% CO_2_. Organoid viability and morphology were assessed after 24 h, and cultures were passaged every 5-7 days using Gentle Cell Dissociation Reagent (STEMCELL Technologies).

### Animals and Animal Models

QTRT1 knockout mice were provided by Dr. Kelly ^31^ and have been previously described ^1^. Animals were provided the same and consistent conditions and utilized in accordance with the UIC Animal Care Committee, the Office of Animal Care and Institutional Biosafety guidelines, and the animal protocol (number ACC 18-179 and ACC 21-120).

QTRT1^LoxP^ mice were generated via the CRISPR/Cas9/Cre method by Cyagen Biosciences (Santa Clara, CA, USA) in the C57BL/6 mouse strain background. The QTRT1 gene (NCBI Reference Sequence: NM_021888.2; Ensembl: ENSMUSG00000002825) is located on mouse chromosome 9 with 10 exons, including the ATG start codon in exon 1 and the TGA stop codon in exon 10 (Transcript: ENSMUST00000002902). Exons 1-3 were selected as the conditional knockout region. The first LoxP was inserted in a non-conservative region (∼1350 bp) upstream of the ATG start codon. The gRNA [gRNA-A1 (matching forward strand of QTRT1 gene): 5′-GTGCTACCATCTTACCTGCCTGG-3′ and gRNA-A2 (matching reverse strand of QTRT1 gene): 5′-ATAAAGAATAGGTCCTGTGTTGG-3′], the donor vector containing LoxP sites, and Cas9 mRNA were co-injected into fertilized mouse eggs for QTRT1^LoxP^ recombinant mice production. The intestinal epithelial cell-specific deletion of QTRT1 mice (QTRT1^ΔIEC^) was achieved by crossing QTRT1^LoxP^ mice with the Villin-Cre mice. The genotyping was performed by using PCR primer pair 5’-AGTATGTGTGCAGAGGACTCAAC-3′ and Reverse 5′-ACGTACCCAGAGGACATGGTA-3′ to identify the LoxP insertion (288 bp for the mutant band and 218 bp for the WT band).

### CRISPR/Cas9-Mediated QTRT1 Knockout in Caco-2 BBE Cells

Caco-2 BBE cells were seeded in 6-well tissue culture plates and maintained in Dulbecco’s modified Eagle’s medium (DMEM) until reaching about 80% confluence. The cells were then transfected with 2 μg of QTRT1 double nickase plasmid (sc-413456-NIC) or control double nickase plasmid (sc-437281) (Santa Cruz Biotechnology, Dallas, TX, USA) using 10 μL of Lipofectamine LTX (Thermo Fisher Scientific, Rockford, IL, USA) and 2.5 μL of PLUS Reagent (Invitrogen, San Diego, CA, USA) per well, according to the manufacturer’s instructions. Transfected cells expressing the GFP marker encoded by the QTRT1 double nickase plasmid were positively selected by flow cytometry using a MoFlo Astrios cell sorter (Beckman Coulter, Indianapolis, IN, USA).

### Treatment of Cells and Human Colonic Organoids

Human colonic organoids derived from IBD patients were established and maintained in growth factor-reduced Matrigel (Corning) and cultured in complete human intestinal organoid medium.^32^ At day 6 after passage, organoids were released from Matrigel using cold basal medium, gently pelleted, and replated in fresh Matrigel domes. After recovery for 24 h, organoids were treated with 10 μM Mitoquinone mesylate (MitoQ; MedChemExpress) diluted in organoid growth medium for 24 h. Control organoids received vehicle treatment (0.1% DMSO). Following treatment, organoids were collected for confocal imaging as indicated.

### Immunoblotting

Mouse intestinal epithelial cells were harvested by gently scraping longitudinally dissected ileum and colon tissues, from proximal to distal end. The cells were collected in lysis buffer (10 mM Tris pH 7.4, 150 mM NaCl, 1 mM EDTA, 1 mM EGTA pH 8.0, 1% Triton X-100, 0.2 mM sodium ortho-vanadate, and protease inhibitor cocktail) and protein concentrations were measured using BioRad Reagent (BioRad), as described previously ^17^. Equal amounts of protein were separated by SDS-polyacrylamide gel electrophoresis, transferred to nitrocellulose membrane, blocked with 5% BSA, and incubated overnight at 4°C with desired primary antibodies. The following day, membranes were washed and incubated with target secondary antibodies for 1 hour and bands were visualized using Enhanced Chemiluminescence detection ^33^ reagent (Thermo Fisher Scientific).

### Histology of mice colon tissues

Mice ileum and proximal colon tissues were harvested, fixed in 10% formalin (pH 7.4), and paraffin-embedded. Slides of tissue sections were prepared at a thickness of 4 μm which were deparaffinized in xylene and rehydrated by passing through graded alcohol. Slides were then stained with hematoxylin and eosin followed by coverslip mounting with permount and were used to determine the histological damage and inflammation scores as previously described ^34^.

### Immunohistochemistry (IHC) staining

After preparation of the 4 μm thick paraffin-embedded ileum and proximal colon sections on slides, antigen retrieval was achieved by incubating the slides for 15 min in hot preheated sodium citrate (pH 6.0) buffer, followed by 30 min of cooling at room temperature. Endogenous peroxidases were quenched by incubating the slides in 3% hydrogen peroxide for 10 min, followed by three rinses with PBS. Slides were then incubated for 1 h with blocking buffer prepared with 2% bovine serum albumin (BSA) (Gemini Bio-products, 700-102P), 1% normal goat serum (Jackson ImmunoResearch Laboratories, 005-000-121), and 1% Triton X-100 (Sigma-Aldrich) in 1X-TBST to reduce nonspecific background. Primary antibodies as listed in Table 1 were applied overnight in a cold room. After three rinses with 1X-TBST, the slides were incubated with secondary antibody (1:100, Jackson ImmunoResearch Laboratories, Cat.No.115-065-174) for 1 h at room temperature. After washing with 1X-TBST for 10 min, the slides were incubated with vectastain ABC reagent (Vector Laboratories, Cat.No. PK-6100) for 1h. After washing with 1X-TBST for five min, color development was achieved by applying a peroxidase substrate kit (Vector Laboratories, Cat.No. SK-4800) for 2 to 5 min, depending on the primary antibody. The duration of peroxidase substrate incubation was determined through pilot experiments and was then held constant for all of the slides. After washing in distilled water, the sections were counterstained with hematoxylin (Leica, Cat.No.3801570), dehydrated through ethanol and xylene, and cover-slipped using a permount (Fisher Scientific, Cat.No. SP15-100). The stained slides were analyzed using the ImageJ Fiji software for Semi-quantification, as described before ^35^.

**Table 1.**
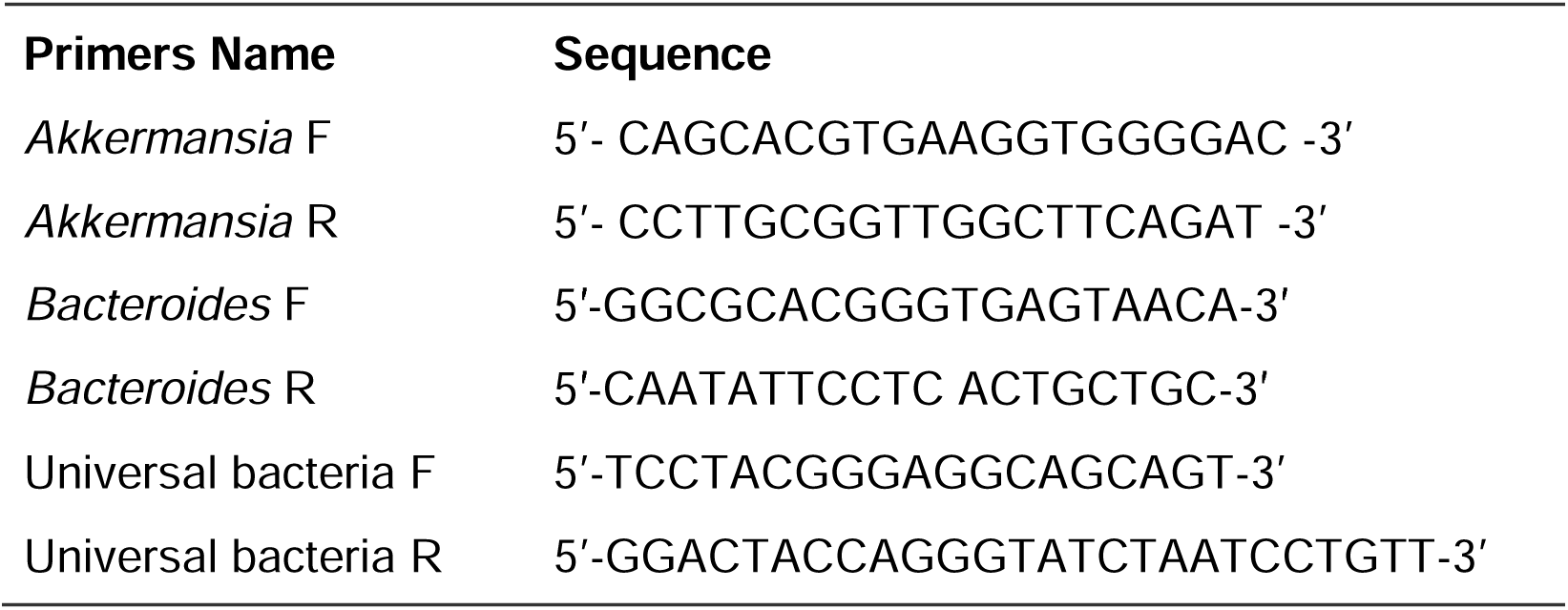
Real-time PCR Primers.

### Immunofluorescence (IF) staining

Mouse ileum and proximal colon tissues were freshly harvested and fixed in 10% neutral buffered formalin followed by paraffin-embedding. Tissue sections were then cut at a thickness of 4μm and prepared on slides for immunofluorescence labelling, as described previously ^36^. Slides were incubated for 1 h with blocking buffer prepared with 2% bovine serum albumin (BSA) (Gemini Bio-products, 700-102P), 1% normal goat serum (Jackson ImmunoResearch Laboratories, 005-000-121), and 1% Triton X-100 (Sigma-Aldrich) in 1X-TBST to reduce nonspecific background. After overnight incubation at 4°C with desired primary antibodies, as listed in Table 1, tissue sections were then labelled with the secondary anti-mouse or anti-rabbit Alexa Fluor 488 and DAPI for 1 hour at room temperature. Lastly, the slides were coverslip mounted with SlowFade (Thermo Fisher Scientific, S2828) and the edges were sealed to prevent from drying. The fluorescence intensity was visualized under a Zeiss laser scanning microscope LSM 710 (Carl Zeiss Inc.) followed by quantitative analysis using ImageJ software.

### MitoTracker-Red Staining and Imaging of human organoids

Human colonic organoids were incubated with MitoTracker™ Red CMXRos (100 nM; Invitrogen, USA) in pre-warmed basal medium for 30 min at 37 °C. Following staining, organoids were gently washed with warm PBS and transferred to glass-bottom dishes for imaging by Zeiss LSM 980 with Airyscan laser scanning microscope equipped with a Z-stack acquisition module. XZ and XY optical sections were captured using identical laser power, gain, and exposure settings for the Normal and IBD groups to allow quantitative comparison. All images were processed uniformly, and no nonlinear adjustments were applied.

### Mitochondrial ROS detection using MitoSOX Red

Mitochondrial superoxide levels were quantified using MitoSOX™ Red mitochondrial superoxide indicator (Invitrogen, USA). Cells were collected and washed twice with warm PBS. MitoSOX Red was added at 5 μM and incubated for 20 min at 37 °C in the dark. After incubation, cells were washed with PBS, resuspended in cold PBS, and immediately analyzed by flow cytometry.

Flow cytometry was performed on a CyTOF Mass Cytometer using excitation at 510 nm and emission at 580 nm. A minimum of 20000 events was collected per sample. Data were processed using FlowJo. MitoSOX-positive cells were gated based on fluorescence intensity using scrambled controls as a baseline.

### Mitochondrial membrane potential measurement using JC-1

Mitochondrial membrane potential (MMP) was assessed using the JC-1 dye (Invitrogen, USA) following manufacturer recommendations. Cells were incubated with 2 μM JC-1 in complete medium for 30 min at 37 °C. After staining, cells were washed with PBS and resuspended in JC-1 assay buffer. Red (J-aggregates) and green (monomers, depolarized mitochondria) fluorescence were recorded on a flow cytometer using 488-nm excitation and 530/590-nm emission filters. Flow cytometric dot plots were divided into quadrants: Q2 (Red-positive) is high membrane potential. Q3 (Green-high) is mitochondrial depolarization. Percentages of JC-1 Red-positive cells were quantified.

### Mitochondrial and Cytosolic Protein Extraction

Mitochondria Isolation Kit for Cultured Cells and Mitochondria Isolation Kit for Tissue were purchased from Thermo Fisher Scientific ^37^. Mitochondrial and cytoplasmic proteins were isolated according to the manufacturer’s instructions with minor modifications. Protein concentrations in mitochondrial and cytoplasmic were determined using a bicinchoninic acid (BCA) protein assay (Thermo Fisher Scientific). Equal amounts of protein from each fraction were used for subsequent SDS-PAGE and Western Blot of mitochondrial proteins and cytoplasmic (released) mitochondrial components.

### Alcian Blue /PAS staining

Alcian blue-PAS staining was performed on paraffin sections of colon tissue fixed with Carnoy’s fixative. Before performing Alcian blue-PAS assay, 5 μm colon tissue sections were baked overnight at 55 °C. Colon tissue sections were deparaffinized in xylene, dehydrated with 100% ethanol. The air-dry slides were stained with Alcian blue solution (1 g of Alcian blue, pH 2.5, 3 mL/L of acetic acid, and 97 mL of distilled water) for 30 min for goblet cells. This was followed by rinsing in tap water for 10 min, oxidizing in periodic acid (5 g/L) for 5 min, rinsing in tap water for 10 min, and staining in Schiff’s reagent as a counter stain (Electron Microscopy Sciences, Cat. NO. 26853-01, Hatfield, PA, USA) for 10 min. After washing in distilled water for 10 minutes, dehydrated through ethanol and xylene, and cover-slipped using a permount (Fisher Scientific, Cat.No. SP15-100, Waltham, MA, USA).

### Intestinal permeability

Fluorescein Dextran (Molecular weight 3 kDa, diluted in HBSS) was gavaged (50 mg/kg mouse). Four hours later, mouse blood samples were collected for fluorescence intensity measurement, as previously reported.^38^

### Fish staining

FISH was performed using antisense single-stranded DNA probes targeting the bacterial 16S ribosomal RNA. The EUB338 ( 5’-GCTGCCTCCCGTAGGAGT-3’) conjugated to Alexa Fluor488 probe was used for universal bacteria. And CY3-AKK (5’-CAGCACGTGAAGGTGGGGAC-3’) was used for *Akkermansia*. FISH was performed on paraffin sections of mucosal biopsies fixed with Carnoy’s fixative or with 10% formalin solution. Before performing the FISH assay, 5 μm tissue sections were baked overnight at 55 °C. Tissue sections were deparaffinized in xylene, dehydrated with 100% ethanol, air dried, incubated in 0.2 mol/L HCl for 20 minutes, and heated in 1 mmol/L sodium thiocyanate at 80 °C for 10 minutes. Samples were pepsin-digested (4% pepsin in 0.01 N HCl) for 20 minutes at 37 °C), slides were washed in wash buffer (0.3 mol/L NaCl, 0.03 mol/L sodium citrate, pH 7, and 0.1% SDS), fixed in 10% buffered formalin for 15 minutes, washed and dried, and hybridized with the probes at 5 ng/ μL concentration each for 5 minutes at 96 °C in hybridization buffer (0.9 mol/L NaCl, 30% formamide, 20 mmol/L Tris-HCl, pH 7.4, and 0.01% SDS) and incubated at 37 °C overnight. Slides were washed 4 times for 5 minutes each at 45 °C in wash buffer. For visualization of the epithelial cell nuclei, the slides were counterstained with 6-diamidino-2-phenylindole/antifade solution. Slides were examined with a Zeiss laser scanning microscope LSM 980 (Carl Zeiss, Inc). Fluorescence staining intensity was determined by using ImageJ software.

### Real-time PCR measurement of bacterial DNA

DNA was extracted from QTRT1^-/-^ and age-matched WT mice fecal samples using EZNA Stool DNA Kit (Omega Bio-tek, Inc. D4015–01, Norcross, CA, USA). The quantitative real-time PCR was conducted using the CFX96 Real-time PCR detection system (Bio-Rad Laboratories, Hercules, CA, USA) and iTaq^TM^ Universal SYBR green supermix (Bio-Rad Laboratories, Hercules, CA, USA) according to the manufacturer’s directions. All expression levels were normalized to universal bacteria levels of the same sample. Percent expression was calculated as the ratio of the normalized value of each sample to that of the corresponding untreated control cells. All real-time PCR reactions were performed in triplicate. Primer sequences were designed using Primer-BLAST or were obtained from Primer Bank primer pairs listed in **Table 1**.

### Statistical analysis

All experiments were repeated at least three times and data represented in bar graphs are expressed as mean ± SD or ± SEM. All performed statistical tests were 2-sided and *P*-values of 0.05 or less were considered statistically significant. Shapiro-Wilk test ^39^ was used to test the normality of data. Comparisons between two groups with normal distribution were analyzed by Welch’s *t*-test or Wilcoxon rank sum test, as appropriate. Comparisons between three or more groups were analyzed by Welch’s ANOVA test based on normal distributions of data. To correct for multiple comparisons in ANOVA, *P*-values were adjusted by Dunnett’s T3 multiple comparison test ^40^. All statistical analyses were performed using GraphPad Prism version 8.0.0 for Windows or latest version of R in Mac.

## Acknowledgements/Funds

We would like to acknowledge the Crohn’s & Colitis Foundation Senior Research Award (Grant No. 902766), UIC Cancer Center, the NIDDK/National Institutes of Health grant R01 DK134343, R01DK114126, VA Merit Award VA 1 I01 BX004824-06, and VA Collaborative Merit Award 1 I01BX006878-01A1 to Jun Sun. The study sponsors play no role in the study design, data collection, analysis, and interpretation of data. The contents do not represent the views of the United States Department of Veterans Affairs or the United States Government. Partial of our data was orally presented at the Digestive Disease Week (DDW) 2025.

## Abbreviation list

AMPs: antimicrobial peptides
CD: Crohn’s disease
DSS: Dextran sulfate sodium
GEO: Gene Expression Omnibus
IBD: Inflammatory bowel disease
IL10: Interleukin 10
KO: knockout
Mito: mitochondria
mtDNA: mitochondrial DNA
MMP: mitochondrial membrane potential
NF-κB: Nuclear factor kappa-light-chain-enhancer of activated B cells
PCNA: Proliferating Cell Nuclear Antigen
Q: queuosine
q: queuine
QTRT1: Queuine tRNA-ribosyltransferase 1
QTRT2: Queuine tRNA-ribosyltransferase accessory subunit 2
Q-tRNA: Queuosine-containing tRNA
ROS: reactive oxygen species
scRNAsq: single cell RNA sequencing
TEER: Transepithelial electrical resistance
TGT: tRNA-guanine transglycosylase
TJ: tight junctions
TNFα: Tumor Necrosis Factor α
UC: Ulcerative colitis
WT: wildtype

## References

1. Zhang J, Zhang Y, McGrenaghan CJ, et al. Disruption to tRNA Modification by Queuine Contributes to Inflammatory Bowel Disease. Cell Mol Gastroenterol Hepatol 2023;15:1371–1389.

2. Fergus C, Barnes D, Alqasem MA, et al. The queuine micronutrient: charting a course from microbe to man. Nutrients 2015;7:2897–929.

3. Diaz-Rullo J, Gonzalez-Pastor JE. tRNA queuosine modification is involved in biofilm formation and virulence in bacteria. Nucleic Acids Res 2023;51:9821–9837.

4. Yan F, Xiang S, Shi L, et al. Synthesis of queuine by colonic gut microbiome via cross-feeding. Food Frontiers 2024;5:174–187.

5. Boland C, Hayes P, Santa-Maria I, et al. Queuosine formation in eukaryotic tRNA occurs via a mitochondria-localized heteromeric transglycosylase. J Biol Chem 2009;284:18218–27.

6. Alula KM, Dowdell AS, LeBere B, et al. Interplay of gut microbiota and host epithelial mitochondrial dysfunction is necessary for the development of spontaneous intestinal inflammation in mice. Microbiome 2023;11:256.

7. Alula KM, Delgado-Deida Y, Callahan R, et al. Inner mitochondrial membrane protein Prohibitin 1 mediates Nix-induced, Parkin-independent mitophagy. Sci Rep 2023;13:18.

8. Haque PS, Kapur N, Barrett TA, et al. Mitochondrial function and gastrointestinal diseases. Nat Rev Gastroenterol Hepatol 2024;21:537–555.

9. Wehkamp J, Wang G, Kubler I, et al. The Paneth cell alpha-defensin deficiency of ileal Crohn’s disease is linked to Wnt/Tcf-4. J Immunol 2007;179:3109–18.

10. Thachil E, Hugot JP, Arbeille B, et al. Abnormal activation of autophagy-induced crinophagy in Paneth cells from patients with Crohn’s disease. Gastroenterology 2012;142:1097–1099 e4.

11. Wu S, Zhang YG, Lu R, et al. Intestinal epithelial vitamin D receptor deletion leads to defective autophagy in colitis. Gut 2015;64:1082–94.

12. Lu R, Zhang Y-g, Xia Y, et al. Paneth Cell Alertness to Pathogens Maintained by Vitamin D Receptors. Gastroenterology 2021;160:1269–1283.

13. Erben U, Loddenkemper C, Doerfel K, et al. A guide to histomorphological evaluation of intestinal inflammation in mouse models. International journal of clinical and experimental pathology 2014;7:4557–76.

14. Mineta K, Yamamoto Y, Yamazaki Y, et al. Predicted expansion of the claudin multigene family. FEBS Lett 2011;585:606–12.

15. Lloyd-Price J, Arze C, Ananthakrishnan AN, et al. Multi-omics of the gut microbial ecosystem in inflammatory bowel diseases. Nature 2019;569:655–662.

16. Tuorto F, Lyko F. Genome recoding by tRNA modifications. Open Biol 2016;6.

17. Zhang YG, Lu R, Wu S, et al. Vitamin D Receptor Protects Against Dysbiosis and Tumorigenesis via the JAK/STAT Pathway in Intestine. Cell Mol Gastroenterol Hepatol 2020;10:729–746.

18. Lu R, Zhang Y, Xia Y, et al. Paneth cell alertness to pathogens maintained by vitamin D receptors. Gastroenterology 2020.

19. Barnett M, Bermingham E, McNabb W, et al. Investigating micronutrients and epigenetic mechanisms in relation to inflammatory bowel disease. Mutat Res 2010;690:71–80.

20. Kinchen J, Chen HH, Parikh K, et al. Structural Remodeling of the Human Colonic Mesenchyme in Inflammatory Bowel Disease. Cell 2018;175:372–386 e17.

21. Lab S. Getting Started with Seurat v5, 2025.

22. Norman JM, Handley SA, Baldridge MT, et al. Disease-specific alterations in the enteric virome in inflammatory bowel disease. Cell 2015;160:447–460.

23. Mancabelli L, Milani C, Lugli GA, et al. Identification of universal gut microbial biomarkers of common human intestinal diseases by meta-analysis. FEMS microbiology ecology 2017;93:fix153.

24. Hall AB, Yassour M, Sauk J, et al. A novel Ruminococcus gnavus clade enriched in inflammatory bowel disease patients. Genome medicine 2017;9:103.

25. Sankarasubramanian J, Ahmad R, Avuthu N, et al. Gut microbiota and metabolic specificity in ulcerative colitis and Crohn’s disease. Frontiers in medicine 2020;7:606298.

26. Pascal V, Pozuelo M, Borruel N, et al. A microbial signature for Crohn’s disease. Gut 2017;66:813–822.

27. Forbes JD, Chen C-y, Knox NC, et al. A comparative study of the gut microbiota in immune-mediated inflammatory diseases—does a common dysbiosis exist? Microbiome 2018;6:221.

28. Franzosa EA, Sirota-Madi A, Avila-Pacheco J, et al. Gut microbiome structure and metabolic activity in inflammatory bowel disease. Nature microbiology 2019;4:293–305.

29. Schirmer M, Franzosa EA, Lloyd-Price J, et al. Dynamics of metatranscription in the inflammatory bowel disease gut microbiome. Nature microbiology 2018;3:337–346.

30. Lloyd-Price J, Arze C, Ananthakrishnan AN, et al. Multi-omics of the gut microbial ecosystem in inflammatory bowel diseases. Nature 2019;569:655–662.

31. Rakovich T, Boland C, Bernstein I, et al. Queuosine deficiency in eukaryotes compromises tyrosine production through increased tetrahydrobiopterin oxidation. J Biol Chem 2011;286:19354–63.

32. Zhang YG, Sun J. Study Bacteria-Host Interactions Using Intestinal Organoids. Methods Mol Biol 2019;1576:249–254.

33. Wirawan E, Vande Walle L, Kersse K, et al. Caspase-mediated cleavage of Beclin-1 inactivates Beclin-1-induced autophagy and enhances apoptosis by promoting the release of proapoptotic factors from mitochondria. Cell Death Dis 2010;1:e18.

34. Wu SP, Yoon S, Zhang YG, et al. Vitamin D receptor pathway is required for probiotic protection in colitis. American Journal of Physiology-Gastrointestinal and Liver Physiology 2015;309:G341–G349.

35. Zhang J, Zhang Y, Xia Y, et al. Imbalance of the intestinal virome and altered viral-bacterial interactions caused by a conditional deletion of the vitamin D receptor. Gut Microbes 2021;13:1957408.

36. Zhang YG, Lu R, Xia Y, et al. Lack of Vitamin D Receptor Leads to Hyperfunction of Claudin-2 in Intestinal Inflammatory Responses. Inflamm Bowel Dis 2019;25:97–110.

37. Tageldin MH, Wallace DB, Gerdes GH, et al. Lumpy skin disease of cattle: an emerging problem in the Sultanate of Oman. Trop Anim Health Prod 2014;46:241–6.

38. Garrett S, Zhang Y, Xia Y, et al. Intestinal Epithelial Axin1 Deficiency Protects Against Colitis via Altered Gut Microbiota. Engineering (Beijing) 2024;35:241–256.

39. Shapiro SS, Wilk MB. An analysis of variance test for normality (complete samples). Biometrika 1965;52:591–611.

40. Xia Y, Sun J. Applied Microbiome Statistics: Correlation, Association, Interaction and Composition. Boca Raton, FL, USA: Chapman and Hall/CRC, 2024.

